# Cell Position-Associated Division Orders and Cell Cycle Durations Shape Asymmetric Trajectories of Cell Fates and Morphological Events in Pre- and Peri-implantation Mouse Embryos

**DOI:** 10.64898/2026.09.24.754028

**Authors:** Huan-huan Yang

## Abstract

A central question in developmental biology is how spatiotemporal embryo morphological events and cell differentiation are precisely coordinated to eventually develop into a mature organism. Previous research has shown cell positions and asymmetric division stages are related to cell lineage specification since morula stage. However, the detailed spatiotemporal dynamics and histories of embryonic cells and their relations with cell lineage specification remain incompletely understood from the onset of embryo development onward. My study on *in-vitro* embryos showed that continuous live cell tracking mapped community-like patterns in cell origin and differentiation. Cell positions, cell temporal factors (such as cell division order and cell cycle) and their histories intricately interacted over pre-and peri-implantation stages. These interactions of spatiotemporal cellular activities directed embryo morphological events and asymmetric cell lineage origin and differentiation throughout 2-cell to around 100-cell stage. In conclusion, this study provides comprehensive insights into the cellular spatiotemporal dynamics of cell inheritance and differentiation during pre-and peri-implantation morphological events and cell lineage specification. By longitudinally integrating spatial and temporal parameters relating to early embryogenesis, the findings not only bridge divergent explanations stemmed from different studies in the field but also offers a refined foundation for evaluating embryo potential and improving outcomes in assisted reproductive technologies, stem cell research, and regenerative medicine.

**Summary:** The earliest stage of embryo development, from the fertilised zygote to the implantation-ready blastocyst, is central not only to understanding embryogenesis, but also to cell totipotency, pluripotency, and the establishment of stem-cell-based models and organoids. After more than two centuries of study, the field has accumulated extensive knowledge of this process, and many researchers now consider its major features largely resolved. In parallel, diverse methodological approaches have yielded multiple theoretical models, some of which have become dominant and appear to foreclose alternative interpretations. This work offers novel insights into pre-and peri-implantation development, from the 2-cell stage through to beyond the 100-cell stage. Because it challenges prevailing models of cell lineage specification during the first three lineages, the conduction, submission of this work and its publication have met with significant delays, sustained obstruction since at least 2022. To prevent further erosion of its novelty and, more importantly, to contribute new knowledge to the field, the work is shared here as a preprint.

## 1. Introduction

In the absence of technological interventions, mammalian development starts with a fertilised single-celled embryo, the zygote. Prior to implantation, the embryo gradually develops into a blastocyst consisting of three primary cell lineages: the trophectoderm (TE), primitive endoderm (PrE) and epiblast (Epi). The latter two have traditionally been thought to segregate from inner cell mass (ICM). Over decades, cell fate decision-making and lineage specification in mammalian embryos have been subject to ongoing debates between the prevailing non-prepatterning views and alternative theories suggesting early pre-patterning (Yang, 2025a).

Multiple studies have reported molecular heterogeneity of embryonic cells at different stages in both human and mouse embryos although traditional theory thought embryonic cells were identical until embryo compaction (Torres-Padilla *et al*., 2007; Płusa *et al*., 2008; Jedrusik *et al*., 2008; Biase *et al*., 2014; Xenopoulos *et al*., 2015; Goolam *et al*., 2016; Junyent *et al*., 2024). My latest work has showed that even cells initiating embryo compaction, hatching, and protrusion-featured PrE migration exhibit spatiotemporal heterogeneity among their seemingly “same” peers or within a lineage both *in vivo* and *in vitro* (Yang, 2025b). This points to a framework where heterogeneous spatiotemporal dynamics of cells guide embryo morphological events and lineage specification. Spatial heterogeneity is well recognised as a crucial cue in cell fate decisions after the 8-cell stage-(Kimber *et al*., 1982; Yang, 2025a). However, the roles of spatial features of embryonic cell individuals and their histories in embryogenesis remain incompletely understood, particularly at earlier stages. Simultaneously, temporal frameworks are vital for embryogenesis (Junyent *et al*., 2024); disruptions in the chronological sequence and pace of development can greatly impact tissue organisation and regional interplay within embryos (Gaunt *et al*., 1988; Qiao *et al*., 2016). Another fundamental question therefore arises: lacking access to external timing mechanisms, how do embryonic cells intrinsically perceive time, establish the order of developmental events, and regulate developmental tempo?

Based on a critical review of existing literature and my multidisciplinary experiences, I hypothesised that spatiotemporal cues of cellular events and their histories play critical roles in both cell fate decisions and morphological events during pre-and peri-implantation stages. Specifically, I aimed to investigate how historical spatial cues (such as changes in cell positions) and timing patterns (specified as cell division order and cell cycle duration) influence cell specification and functions, and ultimately coordinate cell differentiation and community across successive cell generations. To achieve this, I determined to continuously track and systematically analyse *in-vitro* long non-invasive 4D time-lapse and fixed immunostaining imaging of embryos from H2B-GFP mouse line during E1.5 to E4.5.

By reconstructing continuous cell origin and differentiation trees within embryos, my study revealed novel spatiotemporal cues coordinating cooperative clusters and directing heterogenous trajectories across embryonic cells (including mother, daughter, sibling, and cousin cells) over 2-cell to ∼100-cell stage. This work offers valuable insights into the intricate spatiotemporal interaction web during pre-and early peri-implantation mouse development. My findings provide a foundation for future research into the spatiotemporal molecular mechanisms of transgenerational cell inheritance and differentiation during embryo development, stem cell research, and regenerative medicine.

## 2. Results

### 2.1 Establishing Cell Origin Trees via Continuous Longitudinal Cell Tracking and Imaging Matching during Pre-and Early Peri-Implantation Stages

To investigate the spatiotemporal rhythms and dynamics of embryonic cell activities, I established long and short 4D videos of H2B-GFP embryos and tracked individual cells within each embryo. Zygotes were also observed before live imaging. Long 4D videos were thoroughly tracked retrospectively and prospectively. This tracking was cross-referenced with the Sox2-and Gata4-stained Epi and PrE lineages post-videos (Figure 1A, Figure 16A-C); refer to Materials and Methods for methodological details on cell tracking and image matching). I performed three rounds of verification to ensure the accuracy of cell tracking. During the first correction round, two to three corrections of tracked cells were made per embryo between 8-to 32-cell stages, decreasing to zero during the second and third rounds of corrections. After the 32-cell stage, up to the ∼100-cell stage, five to seven corrections of tracked cells were made initially, reducing to two to three, and zero to one during the second and third rounds of corrections. During tracking, 2.3% of 2070 tracked cells across 10 analysed embryos underwent cell death (Figure 1B,C), as identified through the fragmented nuclei morphology visualised by H2B-GFP. These observed cell deaths occurred majorly post-E3.75, roughly in line with the timing when Sox2-positive ICM numbers decreased in (Yang, 2025b). Around 1.4% of 2070 tracked cells were lost during tracking (Figure 1C), mainly due to insufficient H2B-GFP signal (Figure 16B,C). Based on image matching and tracking results, 78%-92.3% of 102 Gata4 cells were tracked, while 75%-88.3% of 112 Sox2 cells were tracked and 99.9% of TE cells were successfully tracked (Figure 1D). This validated cell tracking allowed me to chart cell lineage trees, and map cell dynamic positions, cell division orders and cell cycle dynamics, as well as their relationships spanning pre-and peri-implantation stages (from 2-cell to beyond 100-cell stages). The example trees, each starting with eight branches and their respective developmental histories, were depicted in (Figure 1B).

**Figure 1.**
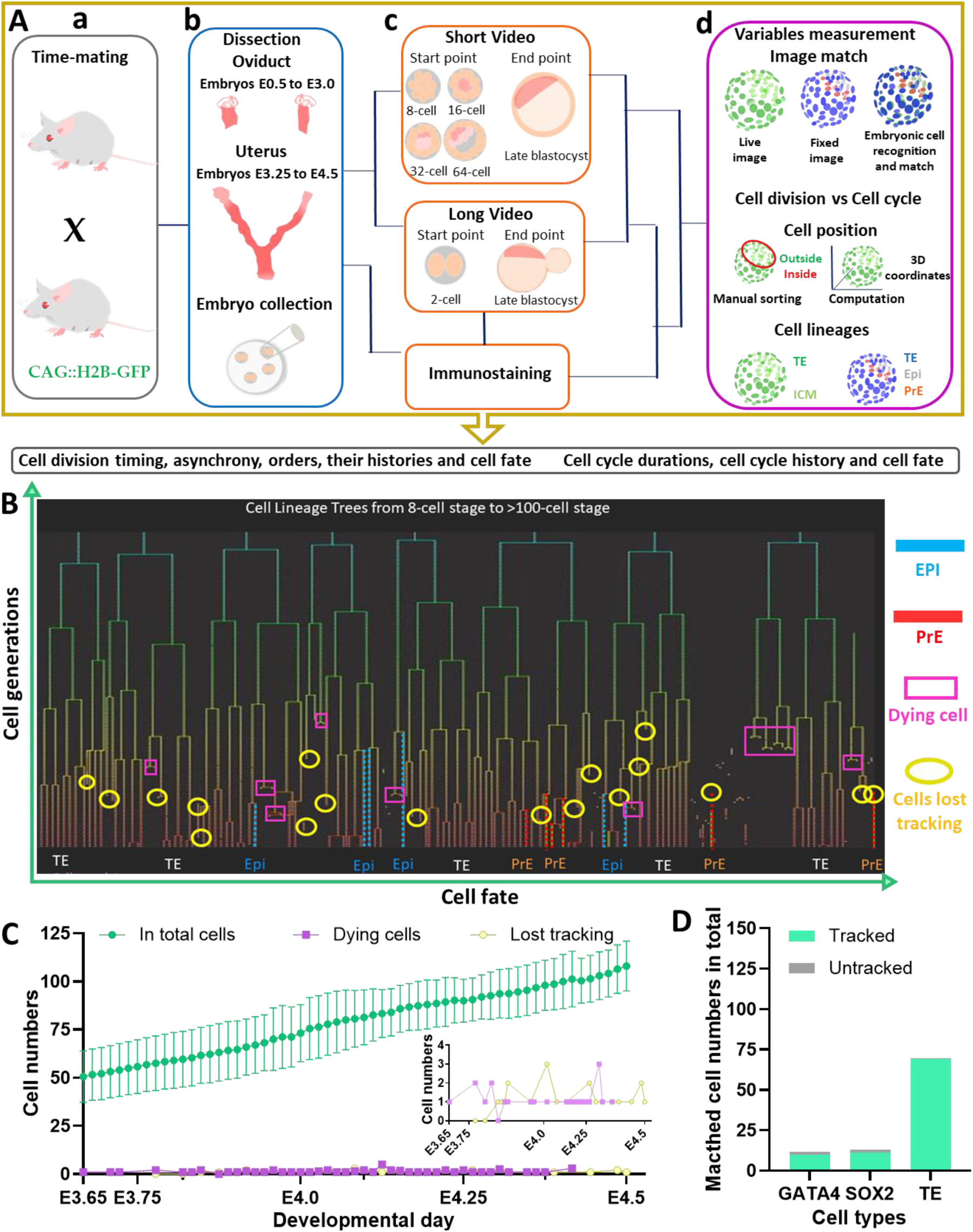
Workflow and cell-lineage tracking trees. **(Aa-c)** Experimental procedures for short (∼36 hours) and long (∼69 hours) videos, with embryos from mice used in (Yang, 2025b). **(Ad)** Image matching methods and major parameter measurements, with result structure outlined (arrow). **(B)** Illustration of successful tracking trees of embryonic cell lineage in an example embryo, with matched cell lineages colour-coded in blue (Sox2-positive cells), red (Gata4-positive cells), yellow (cells losing tracking) and pink (dying cells). **(C)** Total cell numbers (grey) in embryos, cells undergoing death (pink) and cells where continuous tracking was infeasible (yellow) over time. **(D)** Successfully tracked (green) and untracked (grey) identified Sox2, Gata4 and TE cells. Sample size: 7 embryos (from two videos across three litters).

This workflow establishes a validated cell tracking process, ensuring the best possible accuracy of semi-manual cell tracking and matching in this study. The reliable cell lineage tracking trees provide a solid foundation for the analyses detailed in the following sections.

### 2.2 Initial Evaluation of Spatiotemporal Parameters in Long H2B-GFP Embryo Videos

Key spatiotemporal developmental parameters were analysed across mouse litter and among embryos within each litter using long-term *in vitro* imaging. These parameters included the contribution of early embryonic cells to lineages at E4.5, numbers of internalised and externalised cells at the 8-and 16-cell stages, embryonic cell division rhythms and their links with lineage specification, and the relationships between cell cycles and position-specific cell populations (Table 1). Descriptive statistics are presented as (mean with SD) in the table. Man-Whitney test was used to assess inter-litter/video variations (for n ≥ 3 embryos/litter), showing no statistically significant differences in the evaluated parameters, except for general cell cycles in generation 7 (Gn7), and position-specific cycle durations in Gn8 (Table 1). However, these differences were likely due to limited access to all cell cycles of those generations in the live videos before E4.5. Overall, this initiation analysis indicates that embryos filmed for long-term videos from different litters of H2B-GFP mouse line share a consistent baseline of developmental parameters, providing a solid foundation for subsequent analyses.

**Table 1.**
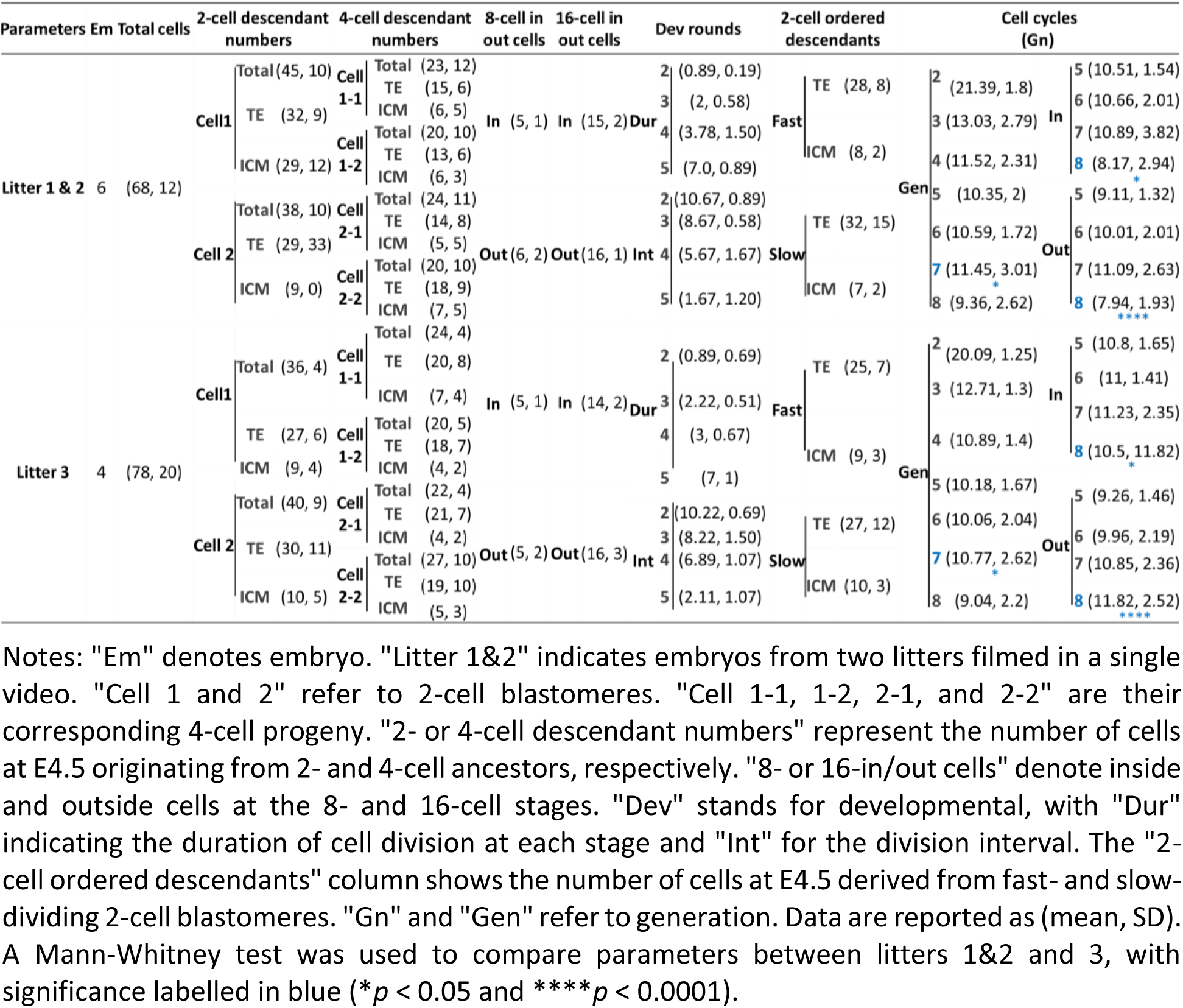
Variability of key analysed parameters across embryos and litters in long H2B-GFP embryo videos.

### 2.3 Historical Positional Cues Driving Lineage Specification in Early Embryogenesis

#### 2.3.1 Mapping cell positions from lineage origins to differentiated cell branches

Using the acquired cell lineage trees (Figure 1B), I mapped the historical positions of each tracked cell during E1.5 and E4.5. Embryonic cells from the same mother consistently tended to cluster together in specific regions over time, regardless of their designation as internal/ICM or external/TE cells. This historical spatial pattern persisted back to the 2-cell stage, with occasional minimal crossover within cell generations from different mother cells (Figure 2A). During the 2-to ∼31-cell stages, embryos and cells underwent detectable rotating displacement within the ZP, especially during cell divisions. Despite these global and local movements, including cell internalisation during the transition from 8 to 16 cells and 16 to 31/32 cells, the positions of cell generations from the same mothers remained distributed within similar regions (Figure 2A).

**Figure 2.**
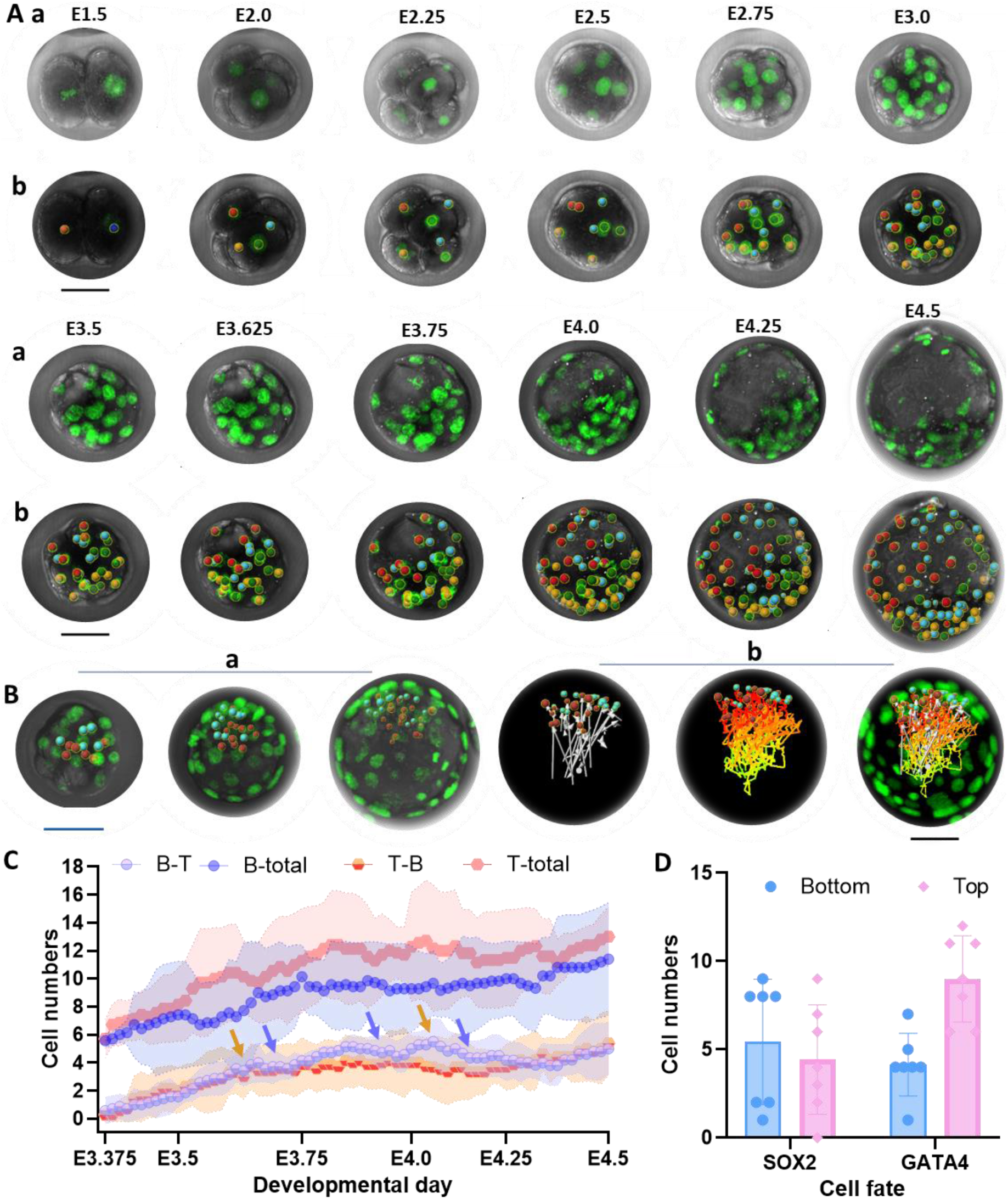
Historical patterns of origin and development of cell position and cell lineages. **(Aa,b)** Embryo development from two cells to over 100 cells (E1.5-E4.5) and their origin, movement and destination, with red and yellow spots representing descendants from the same 2-cell blastomeres (orange at E1.5), blue and green stands for cells sharing the other 2-cell mothers (dark blue at E1.5). **(Ba,b)** Movement and destination of top ICM cells (red spots) and bottom ICM cells (blue spots) over cavity expansion, with ICM cell displacement directions (white arrows) and movement paths (Bb). Scale bar in (A-C): 40 µm. **(C)** Relative movement of top and bottom ICM descendants over E3.75 and E4.5, indicated by cell numbers with SD error bars, including total cell number of the bottom (dark blue) and top (pink) ICM descendants, numbers of bottom cells moving to the top (purple) and top cells moving to the bottom (red/pink). Orange and blue arrows mark notable cell numbers decrease. **(D)** Numbers of the bottom (blue) and top (pink) ICM cells specified into the Epi (Sox2) and PrE (Gata4) lineages. Wilcoxon matched-pairs signed rank test was applied. Error bar: SD. Embryo sample sizes: 7 (from two videos across three litters) in (A-D).

Starting from the 32-cell stage, embryos and their cells ceased rotation since the initiation of cavitation. During cavitation, ICM cells notably shifted to one side within embryos, yet descendants from the same 2-, 4-or 8-cell ancestors continued to cluster in adjacent regions. This clustering applied to TE and ICM cells from the same mothers, although the cavity separated mural TE (Mu-TE) and ICM into abembryonic and embryonic regions (Figure 2A). After the 32-cell stage till the late blastocyst, dynamic movement within ICM was observed. To investigate this, I categorised ICM cells into the bottom (not facing the cavity) and top (facing the cavity) groups 60-80 minutes post-cavity initiation (Figure 2Ba,b). Between the 32-and 64-cell stages, 37.5%-67% of six to 16 bottom ICM cells per embryo moved to the ICM surface, while 10%-50% of eight to 19 top ICM cells moved to the lower ICM (Figure 4.2, Panel C). After E3.75 (∼64 cells), ICM movements between the top and bottom appeared to slow down, with 0%-29% of seven to 17 ICM bottom cells moving to the top, and 0.07% to 20% of 10 to 16 top ICM cells moving to the bottom (Figure 2C). Despite these movements, origin-specific regional clustering persisted over time in 3D viewers. At E4.5, bottom ICM descendants tended to form slightly more Sox2 cells (5 ± 3) and fewer Gata4 cells (4 ± 2) than top ICM descendants, which formed significantly fewer Sox2 cells (4 ± 3) and more Gata4 cells (9 ± 2) (Wilcoxon matched-pairs signed rank test, *p* = 0.035) (Figure 2D).

These data show that the spatial organisation of embryonic cells over time maintains origin-specific regional clustering and influences lineage allocation, with the bottom ICM favouring Sox2-positive cells and the top ICM favouring Gata4-positive cells. These findings underscore the crucial role of spatial dynamics in lineage specification.

#### 2.3.2 Historical spatial cues in cell origin and differentiation at the 2-and 4-cell stages

The mapped dynamic cell positions allowed me to track cell lineage origin and differentiation histories back to the 2-and 4-cell stages. The results showed asymmetric contributions of 2-cell blastomeres to total embryonic cells, TE and ICM lineages, with a relative switch in TE contributions between E3.5 and E3.75-E4.5 (Figure 3Aa-d). At E4.5, in nine of 10 embryos, progeny from one 2-cell blastomere (termed cell A) constituted 50%-100% of four to 15 Gata4 cells and 0% to 50% of two to 14 Sox2 cells per embryo (Figure 3Ad). The other 2-cell blastomere (cell B) contributed to 50%-100% Sox2 cells and 0%-50% Gata4 cells (Figure 3Ad). Such asymmetric contributions of 2-cell blastomeres differed significantly (Wilcoxon matched-pairs signed rank test, *p* = 0.027 for Sox2 cells, *p* = 0.006 for Gata4 cells) (Figure 3Ab-d). To determine factors influencing asymmetric cell fate allocation, I analysed the distance between cell nuclei and the nearest free-contact cell surface (nuclear-surface distance). Cells at the 2-cell stage, generating more Gata4 progeny, showed significantly longer nuclear-surface distances (24.7 ± 2.3 µm) than the cells generating more Sox2 cells (22.2 ± 2.8 µm) (Wilcoxon matched-pairs signed rank test, *p* = 0.001) (Figure 4.3, Panel B).

Similarly, the 4-cell blastomeres tended to asymmetrically contribute to total embryo cells, TE and ICM lineages at E4.5. Across 60% of 10 embryos, 4-cell blastomeres (termed cells A1-1) originating from 2-cell blastomere A consistently formed more polar TE (P-TE) cells and more Gata4 cells than their pairwise sisters (cell A1-2) (Figure 3Ca-d). Similar trends were seen between the other pairwise 4-cell blastomeres (cells B-1 and B-2) from 2-cell blastomeres B per embryo across 70% of embryos. I then analysed 4-cell blastomere positions by measuring their nuclei distances from embryo geometric centres. The result showed that in 80% of 10 embryos, A-1 cells producing more Gata4 cells were positioned further from the embryo centre than their pairwise A-2 sisters. In the remaining 20%, where A-1 blastomeres were closer to embryo centres, A-2 cells with further positions contributed equally to Gata4 and Sox2 cells per embryo (Figure 3D). In 60% of embryos, B-1 cells generating more Gata4 cells were further from embryo centres. In the other 40%, B-2 cells were relatively further than their pairwise B-1 sisters; in half of these cases, B-2 cells produced more Gata4 cells, while in the other half, they generated the equal number of Gata4 and Sox2 cells (Figure 3D). Furthermore, migrating pioneering and hatching breaker cells shared ancestry tracked back to the 2-, 4-and 8-cell stages in 50%, 30%, and 20% of 10 embryos, with their 2-cell mother forming more Gata4 cells.

**Figure 3.**
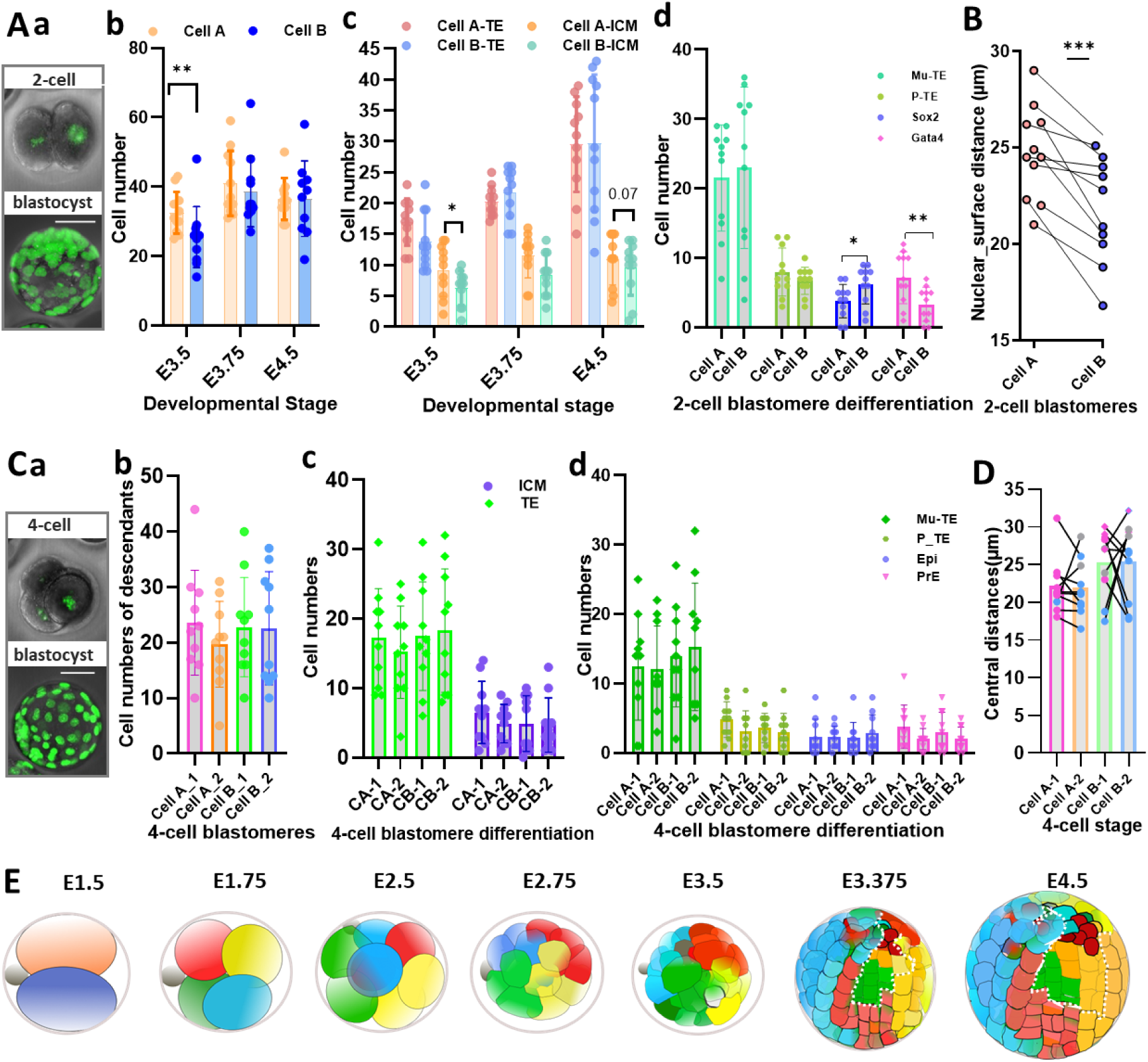
Asymmetric differentiation history of 2-cell and 4-cell blastomeres. **(Aa)** Embryo development from E1.5 to E4.5. **(Ab)** Total cell numbers originating from each 2-cell blastomere. **(Ac)** Cell numbers of TE and ICM originating from 2-cell blastomeres. **(Ad)** Cell numbers of various lineages from the 2-cell blastomeres, including Mu-TE (light green), P-TE (dark blue), Epi (light blue) and PrE (pink). **(B)** Nuclei distance to the nearest free cell surface, with plots in the same colour for the 2-cell blastomere pairs (cell A and cell B). **(Ca)** Embryo development from 4-cell stage to late blastocyst. **(Cb)** Total cell numbers originating from each 4-cell blastomere. Panel **(Cc)** Cell numbers of TE and ICM from each 4-cell blastomere. Panel Cd: Cell numbers of Mu-TE, P-TE, Epi and PrE from each 4-cell blastomere. **(D)** Cell positions impacting 4-cell blastomere asymmetrical development. Positions were calculated via the distances between each cell nucleus and embryonic geometrical centres. Colour-coded points refer to 4-cell blastomere ancestors producing the PrE and Epi: pink for Gata4, blue for Sox2, and grey for cells contributing to equal numbers of Epi and PrE. **(E)** Mapping of cell origin and differentiation during pre-and peri-implantation, with descendants in red and yellow from the same 2-cell mother (orange), blue and green from the other 2-cell mother (dark blue). Cells with black contours indicate ICM. The inside areas circled by white dash show the internal view of embryos. Scale bar in Panel Aa, Ba, Ca, and Da: 40 µm. The error bar denotes SD. In (A,B), statistical differences were determined by Wilcoxon matched-pairs signed rank test, with \**p* < 0.05, \*\**p* < 0.01, and \*\*\**p* < 0.001. Sample size: 11 embryos (from two videos across three litters) in (A-E).

These data reveal that 2-and 4-cell blastomeres exhibit distinct tendencies to form ICM, TE, Epi and PrE, driven by cell position histories. These findings highlight the links between early blastomere positions and cell fate decisions in early embryogenesis.

#### 2.3.3 Historical spatial cues in cell differentiation at the morula stages

The 8-cell blastomeres showed asymmetrical developmental trajectories, particularly across pairwise sisters, seen as early as cell internalisation and externalisation (Figure 4Aa-c). In each embryo (n=10), notably more 8-cell mothers (three to seven) internalised one daughter during the 8-to 16-cell transition, while fewer mothers (one to five) produced both daughters staying outside (Wilcoxon matched-pairs signed rank test, *p* = 0.078) (Figure 4Ab,Ac, yellow and blue lines respectively). At the 16-cell stage, external and internal cells per embryo were 10 to 13 and three to six, respectively. During 16-to 32-cell transition, two to 10 external 16-cell blastomeres per embryo, originating from the 8-cell ancestors without internalising daughters during the 8-to 16-cell transition, internalised one or both of daughters (one out of 10 embryos). In three of 10 embryos, one to three 8-cell blastomeres internalised zero daughters between 8-to 32-cell stages. Zero to two 16-cell blastomeres, derived from external 8-cell blastomeres with one daughter already internalised during the 8-to 16-cell stage, internalised one daughter during the 16-to 32-transition. These asymmetrical internalisations between 16-cell external blastomeres originating from distinct 8-cell blastomeres differed significantly (Wilcoxon matched-pairs signed rank test, *p* = 0.043) (Figure 4Ab,Ac, pink and green respectively). During the 16-to 32-cell stage, when both daughters of a blastomere internalised, one was later externalised. During the 32-to 64-cell transition, one to two cells were internalised in three of 10 embryos. At the 32-cell stage, each embryo had 15 to 17 cells in both external and internal groups, with three to six and three to ten cells newly internalised during 8-to 16-cell and 16-to 32-cell transitions.

**Figure 4.**
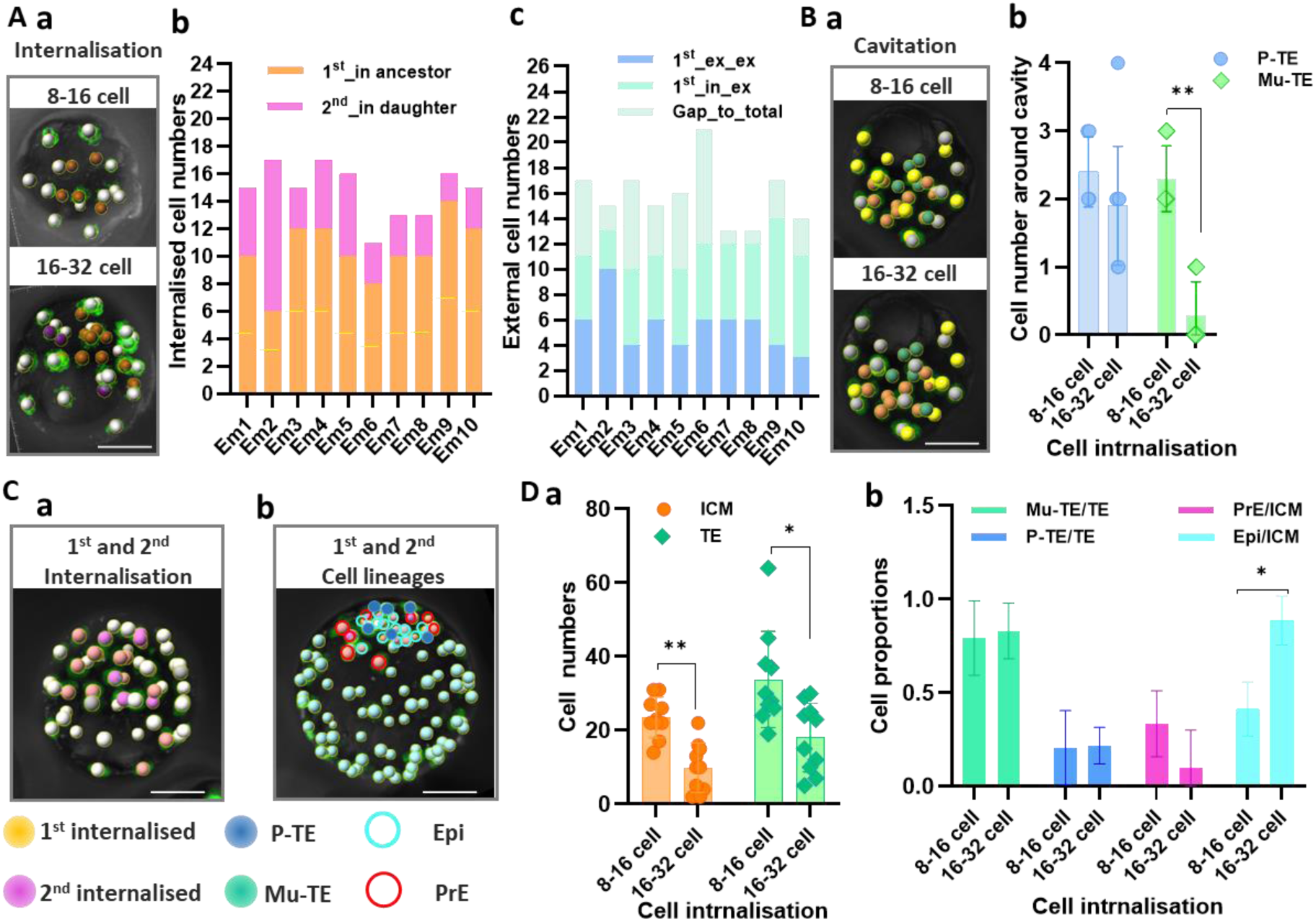
Asymmetric developmental trajectories of 8-cell and 16-cell blastomeres. **(Aa)** Internalised cells during the 8-to 16-cell transition (top) and 16-to 32-cell (bottom) transitions. **(Ab)** Origin of internalised cells at 32-cell stages, including the internalised ICM cell from the 8-to 16-cell stage (at yellow line in orange) and their daughters from the 16-to 32-cell stage (above the yellow line), internalised cells during the 16-to 32-cell transition (pink). **(Ac)** Numbers of TE cells at the beginning of the 32-cell stage and their origin, including external cells (blue) remaining outside from the 8-to 16-and to 32-cell stages, external cells (green) internalising daughters during the 16-to 32-cell stage, external cells (light green) remaining outside after cell internalisation during 16-to 32-cell stage. “1^st^” and “2^nd^” refer to 8-to 16-cell internalisation and 16-to 32-cell internalisation, respectively. **(Ba,b)** Numbers of TE cells, surrounding the cavity initiation, sharing the origin with the 1^st^ or the 2^nd^ internalised ICM cells. “P” and “Mu” refer to the P-TE (blue) and Mu-TE (green) cells, respectively. **(Ca,b)** Sequential allocation of the 1^st^ (orange spots) and 2^nd^ (pink spots) internalised cells and cell lineages after image match, including P–TE (dark blue), Mu-TE (green), Epi (light blue) and PrE (red). **(Da,b)** Cell numbers of ICM (orange) and TE (green) lineages at E4.5, with Mu-TE (green), P-TE (blue), Epi (light blue) and PrE (pink) lineages originating from ancestors internalised daughter cells at the 8-to 16-cell and 16-to 32-cell stages, respectively. Scale bar in (Aa,Ba,Ca and Cb): 40µm. Embryo sample sizes: 10 (from two videos across three litters) in Panels Bb, Da and Db. Error bar: SD, and \**p* < 0.05, \*\**p* < 0.01.

The historical relations between internalised cells and cavity initiation showed that significantly more TE cells sharing the origin with the 1^st^ internalised cells clustered in the Mu-TE region around the initiated cavity than the TE cells sharing the origin with the 2^nd^ internalised cells (Wilcoxon matched-pairs signed rank test, *p* = 0.002) (Figure 4Ba,b). The relations between internalised cell ancestors and cell fates displayed that ancestor cells producing internalised cells during the 8-to 16-cell stage contributed to more ICM (24 ± 5) and more TE (34 ± 13) cells than the 8-cell ancestors generating cells internalised during the 16-to 32-cell stage (10 ± 6 ICM cells vs 18 ± 9 TE cells) (Wilcoxon matched-pairs signed rank test, *p* = 0.002 and *p* = 0.010, respectively) (Figure 4Ca,b;Da,b). Furthermore, the 1^st^ internalised ICM tended to form higher proportions of PrE while the 2^nd^ internalised ICM developed into significantly higher proportions of Epi (Wilcoxon matched-pairs signed rank test, *p* = 0.016) (Figure 4Db).

These data show that cells, producing more internalised daughters in the 1^st^ internalisation round, cluster as Mu-TE around the initiated cavity and also contribute to more PrE cells. These findings provide new insights into the relationships between spatiotemporal internalisation and cell lineage allocation.

### 2.4 Cell Division Order-Associated Cell Lineage Specification in Early Mouse Embryo

My data analysis showed specific 24-hour time patterns of embryonic cell divisions throughout pre-and peri-implantation stages (refer to Appendix A, Figure A1 for further details). It provided the foundation for the detailed investigation into division rates of individual cells within embryos and their relationships with other embryonic cellular features/events, such as cell lineage specification.

#### 2.4.1 Temporal dynamics of embryonic cell division at pre-and peri-implantation stages

To explore temporal dynamics in early embryogenesis, I analysed individual cell division timings via long 4D-filmed H2B-GFP embryos. Cell divisions at each stage were not instantaneous per se but occurred over notable continuation periods, followed by non-division intervals (Figure 5A). To consistently describe this, I developed a conceptual framework for the “embryonic progression period”, “embryonic division (round) durations”, “embryonic division intervals” and their links with cell generations and cell number-specified stages (Figure 5B; refer to Appendix B, C for the details of aforesaid conceptions). Division duration in each round was measured from the moment when embryos first exited the previous stage characterised by cell number 2^n^ (n for division round and n ∈ Z^+^) until all embryonic cells from the 2^n^-cell stage had completed their divisions and embryos completely entered into the 2^(n+1)^-cell stage. Division interval was calculated from the moment when embryos completely entered the 2^(n+1)^-cell stage to the time when embryos exited this stage and started to enter the next 2^(n+2)^-cell stage. Embryonic division durations increased from 0.89 ± 0.46, 2.11 ± 0.50, 3.39 ± 1.42 to 7 ± 0.84 hours at the 2^nd^, 3^rd^, 4^th^ and 5^th^ division rounds, respectively, between E1.5 to ∼E4.5 (six embryos). Division intervals correspondingly decreased from 10.44 ± 0.75, 8.44 ± 1.05, 6.28 ± 1.42 to 1.89 ± 1.05 hours (Figure 5C-D). As embryos reached ∼50 cells, cell division timings overlapped between two successive rounds, without clear division intervals. Between the 5^th^-6^th^ rounds, 2 ± 1 P-TE cells near Mu-TE started the 6^th^ division round, while one to two P-TE and ICM cells had not yet started their 5^th^ round (Figure 5 C,E,F). Between the 6^th^ and 7^th^ division rounds, one to two P-TE cells near Mu-TE started the 7^th^ round, while 3 ± 2 P-TE and ICM cells had not yet started their 6^th^ round. Between the 7^th^ and 8^th^ rounds, 2 ± 1 P-TE (including those near Mu-TE) started their 8^th^ round but 8 ± 7 Mu-TE cells had not yet started their 7^th^ division rounds.

**Figure 5.**
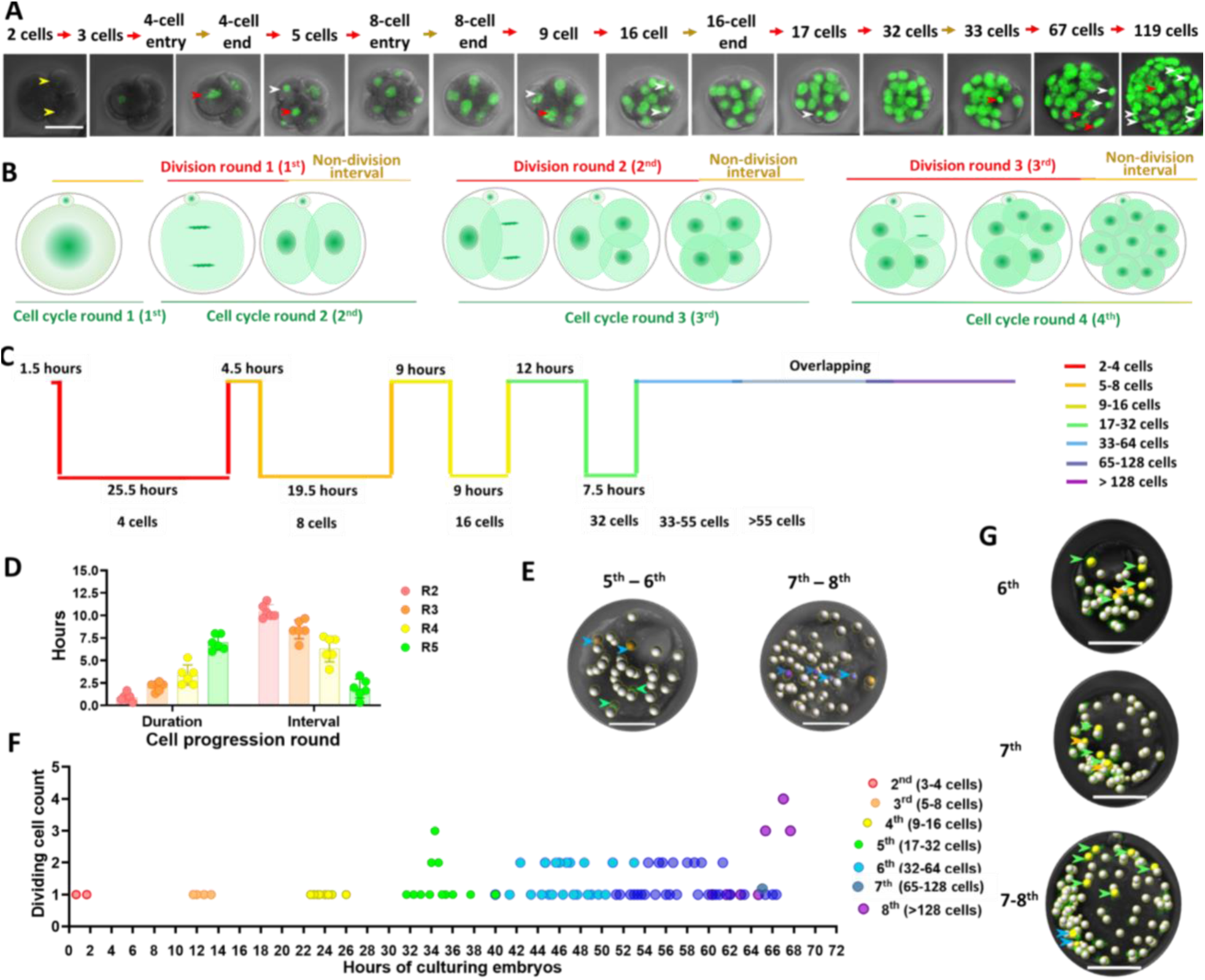
Embryonic cell division dynamics during pre-and peri-implantation. **(A)** Embryonic cell cycle progression of entrance and exit of different developmental stages, including timing with cell divisions (red directional arrow) and without cell divisions (yellow directional arrow). White and red arrows within embryos show cells that just divided and were about to divide, respectively. **(B)** Conception framework of embryonic cell progression round (green lines), embryonic cell division round/duration (red lines) and non-division interval (yellow lines). **(C)** Embryonic cell division round duration and interval lengths in the representative embryo in the 2^nd^ (red), 3^rd^ (orange), 4^th^ (yellow), 5^th^ (green), 6^th^ (blue), 7^th^ (dark blue), and 8^th^ (purple) rounds. **(D)** Distribution of division durations and intervals with SD error bar across six embryos from progression rounds 2 to 5. **(E)** Positions of dividing cells overlapped between two continuous division rounds (5^th^-6^th^, 7^th^-8^th^), showing the latest dividing cell (green arrow) in the 5^th^ round and the earliest dividing cell (blue) in the 6^th^ round, the latest dividing cell (blue arrow) in the 6^th^ round and the earliest dividing cell (dark blue) in the 7^th^ round. **(F)** Embryonic cell count dividing over time at each division round (in one representative embryo), with each round colour-coded as shown in Panel C. **(G)** Synchronously dividing cells dividing in the late division rounds across P-TE (green), ICM (orange) green, and P-TE (blue). Sample size: six embryos from three litters. The scale bar in Panels A, D and E: 40 µm.

Embryonic division round duration reflected asynchronous cell division (Figure 5A). To examine changes in asynchrony or synchrony (determined using 15-minute intervals based on long 4D video time settings) over division rounds, I investigated dividing cell counts over time in each division round. The result showed that asynchronous divisions predominated over time (Figure 5F), as previously reported (Kelly *et al*., 1978). By the 5^th^ division round, synchronous division emerged, with two to three cells dividing synchronously in the 5^th^ - 7^th^ rounds and 4 ± 2 cells dividing synchronously in the 8^th^ round (Figure 5F). Cells dividing synchronously in the 5^th^ and 6^th^ rounds were P-TE cells and nearby deep or surface ICM cells sharing the same origin in the early 7^th^ round, Mu-TE in the late 7^th^ round, and P-TE in the early 8^th^ round (Figure 5G).

These data demonstrate that cell division durations increase while intervals corresponding decrease over each progression stage/period, with asynchronous divisions dominating and local synchronous divisions gradually emerging. These findings highlight the evolving temporal dynamics of cell divisions during early embryo development.

#### 2.4.2 Relations between cell division orders and cell lineage specification

Since the asynchronous division was the predominant pattern in each progression round, I investigated the historical relationships of asynchronous division orders and cell lineages. It should be noted that division orders and sequences were interchanged in this study. Cell types at the end of the long videos (E4.5) were categorised as shown in (Yang, 2025b Section 3.5.2). My data showed that the descendants of the early-dividing 2-cell blastomeres consistently produced less Epi (3 ± 2) but more PrE (6 ± 3 cells) compared to the descendants of the later-dividing 2-cell ancestors (6 ± 2 cells for Epi and 3 ± 2 cells for PrE) (Figure 6A-D). TE cells from both ancestors during the 2^nd^ division round showed no clear trend of cell counts at E4.5 (Figure 6B).

**Figure 6.**
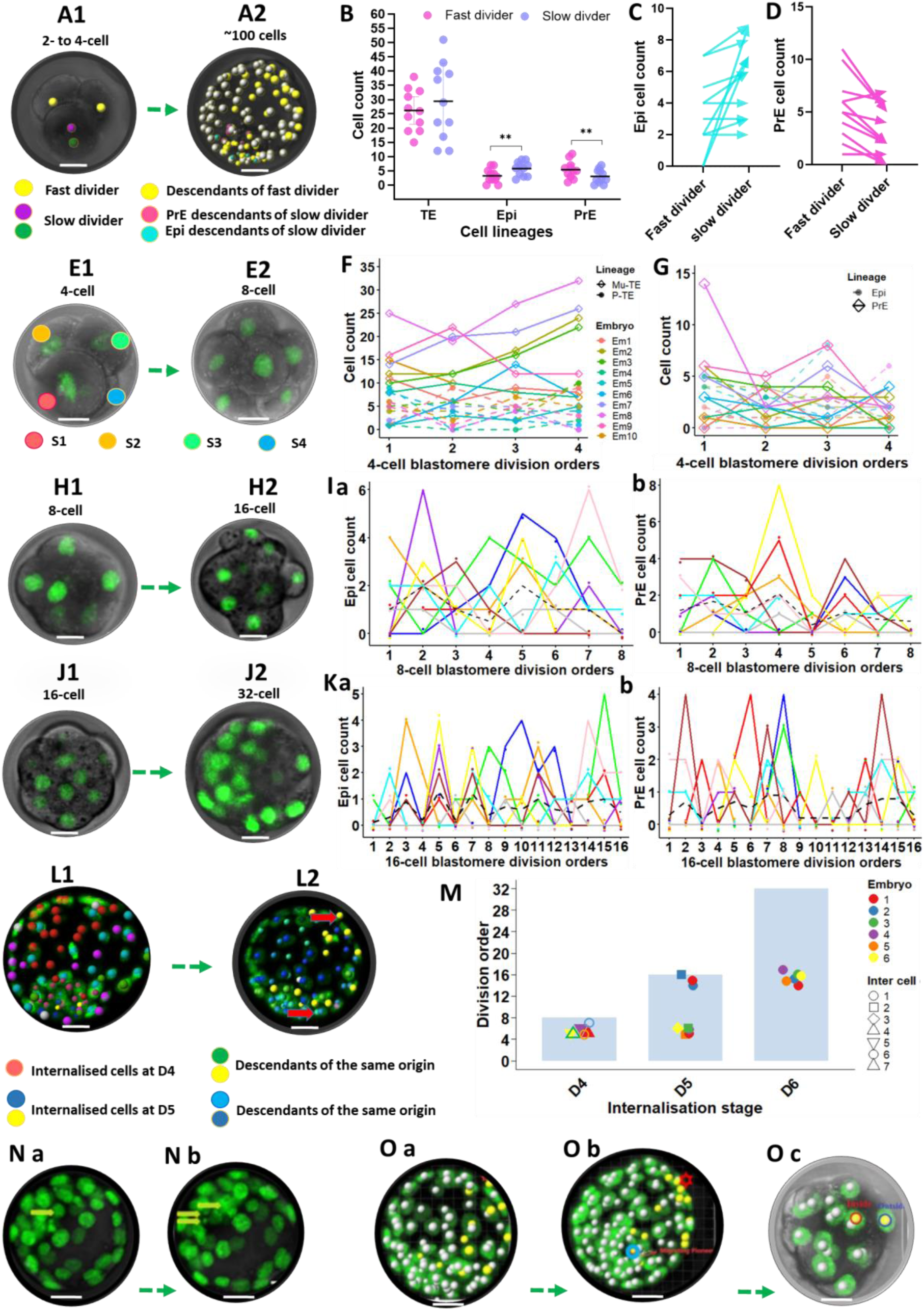
The relations of cell division sequences and cell fate decision. **(A)** Relationships between 2-cell ancestor division orders and distinct cell lineages. **(B-D)** Relationships between 2-cell division orders and cell lineages (Epi in light blue, PrE in pink) and Wilcoxon matched-pairs signed rank test results. **(E-G)** Relations between cell division orders of 4-cell stage and cell lineage specification of P-TE (dashed line), Mu-TE (solid line), Epi (round), and PrE (rhombus). **(H,I)** Relationships between 8-cell division orders and ICM cell lineages (Epi and PrE). **(J,K)** Same as Panels H and I but for 16-cell division orders. **(L,M)** Relations between cell internalisation round and division order of cell lineages, with D for division rounds. Point shapes denote different newly internalised cells differentiating into PrE (Inter cell) in each round, with shapes 6 and 7 shown as unfilled. The same colour of points presents cells from the same embryo. Blue area represents cells other than the newly internalised PrE-committed cells. **(N)** Cells internalised in the 6^th^ round experiencing cell death. **(O)** Relationships between migrating pioneer cells and cell internalisation rounds. In (F,G,I,J,K,M), each colour-coded line represents each embryo. Embryo sample size: 10 (from two videos across three litters). Error bars: SD. The scale bar: 20 µm.

The descendants of late-dividing 4-cell blastomeres in the 3^rd^ round tended to produce more Mu-TE cells: cells in orders 1, 2, 3, and 4 produced 11 ± 7, 12 ± 6, 13 ± 6, and 15 ± 9 Mu-TE cells, respectively. This trend kept relatively consistent in 70% of 10 embryos (Figure 6E,F). The 4-cell ancestors with odd-numbered division orders generated 5 ± 2 and 4 ± 2 P-TE cells near Mu-TE, while those with even-numbered sequences formed 3 ± 3 and 3 ± 2 P-TE cells near Mu-TE (Figure 6F). Furthermore, the descendants of the earliest-dividing ancestors contributed 19%-67% (25^th^ and 75^th^ percentiles) ICM cells in each embryo, with this contribution decreasing to 8%-33%, 10%-38% and 5%-36% for descendants of 4-cell ancestors dividing in orders 2, 3 and 4, respectively (Figure 6G). Epi cells from 4-cell ancestors with even-numbered (2, 4) division orders, appearing in 60% of 10 embryos, outweighed those from ancestors with odd-numbered (1, 3) orders, which occurred in 40% of cases (Figure 6G). PrE cells were more frequently from 4-cell ancestors with the even-numbered (1, 3), appearing in 80% of 10 embryos, than those from ancestors with the odd-numbered, appearing in 20% of cases (Figure 6G).

The descendants of odd-and even-numbered 8-cell ancestors contributed to more Epi and more PrE, respectively. Within each embryo, Epi decedents mainly originated from a single odd-numbered dividing 8-cell ancestor (Figure 6Ia, each peak), with additional contributions from the 8-cell ancestors with division orders closest to the single peak odd-numbered order (Figure 6Ia). In contrast, within each embryo, PrE descendants predominantly arose from two even-numbered dividing 8-cell ancestors (Figure 6Ib). Across 70% of 10 analysed embryos, Epi cells per embryo originated from two to four or zero to three 16-cell blastomeres with odd-and even-numbered division orders, respectively (Figure 6J,Ka). Only 10% of embryos had equal counts of cells with odd-and even-numbered orders forming Epi. In 20% of embryos, two to three odd-division and three to five even-division cells developed into Epi (Figure 6Ka). In total, odd-and even-division 16-cell ancestors gave rise to 50%-89% and 12.5%-50% (25^th^ and 75% percentiles) Epi cells, respectively (Figure 6Ka). In contrast, 50%, 30%, and 20% of embryos had two to four vs zero to two, equal, two to three vs five cells with odd-vs even-division orders forming PrE cells. Within per embryo, 41%-64% and 36%-58% (25^th^ and 75% percentiles) of PrE cells originated from odd-and even-division ancestors, respectively (Figure 6Kb). Notably, 16-cell blastomeres with the earliest order and latest orders tended to rarely contribute to ICM, with cells from 9-11 numbered division orders infrequently contributing to PrE per embryo (Figure 6K).

To further investigate differences between PrE cells internalising at the 4^th^ and the 5^th^ rounds, I tracked the first one to three migrating cells—determined by cell nuclei movement and cell protrusions (Yang, 2025b), and analysed their origins. These detectable migrating pioneer cells seemed to originate from mid-to late-dividing 8-cell ancestors (orders 5 to 7 per embryo) and from relatively early-(orders four to six) or late-dividing (14 to 16) 16-cell ancestors (Figure 6L,M). Additionally, I noticed that very few mid-dividing cells (orders 10 to 13), three cells within the four observed embryos, were internalised in the 6^th^ round (Figure 6M). These internalised cells showed the earliest division orders within ICM cells during the 5^th^ and 6^th^ rounds (orders 1-3 in the 5^th^ round and order 1 in the 6^th^ round) (Figure 6M). Furthermore, cells internalised in the 6^th^ round consistently showed cell protrusions but they more likely died soon after the internalisation (Figure 6N-O).

These data reveal that odd-and even-numbered dividing 2-and 4-cell blastomeres tend to contribute primarily to PrE and Epi cells, respectively, contrasting with the trend observed in 8-and 16-cell blastomeres. Furthermore, my data show PrE cells originate from 8-and 16-cell ancestors with divisions mainly clustered in relatively earlier orders and a few cells dividing in mid-late orders contribute to migrating PrE. These findings suggest the relationships between cell division orders and cell lineage allocation, providing critical insights into the influence of division temporal sequences in embryo development.

#### 2.4.3 Regulators of the relationships between cell division orders and cell lineages

I then examined the influence of cell histories, origins and current positions on the observed non-linear alternating relationships between cell division orders and lineages (Figure 7A; see Materials and Methods for detailed analytical approaches). All mothers were arranged in division orders (Figure 7A-E). Sequencing division orders of each blastomere per generation showed that 4-, 8-, and 16-cell blastomeres, dividing from earliest to latest orders, tended to be progressively closer to embryo centre.

**Figure 7.**
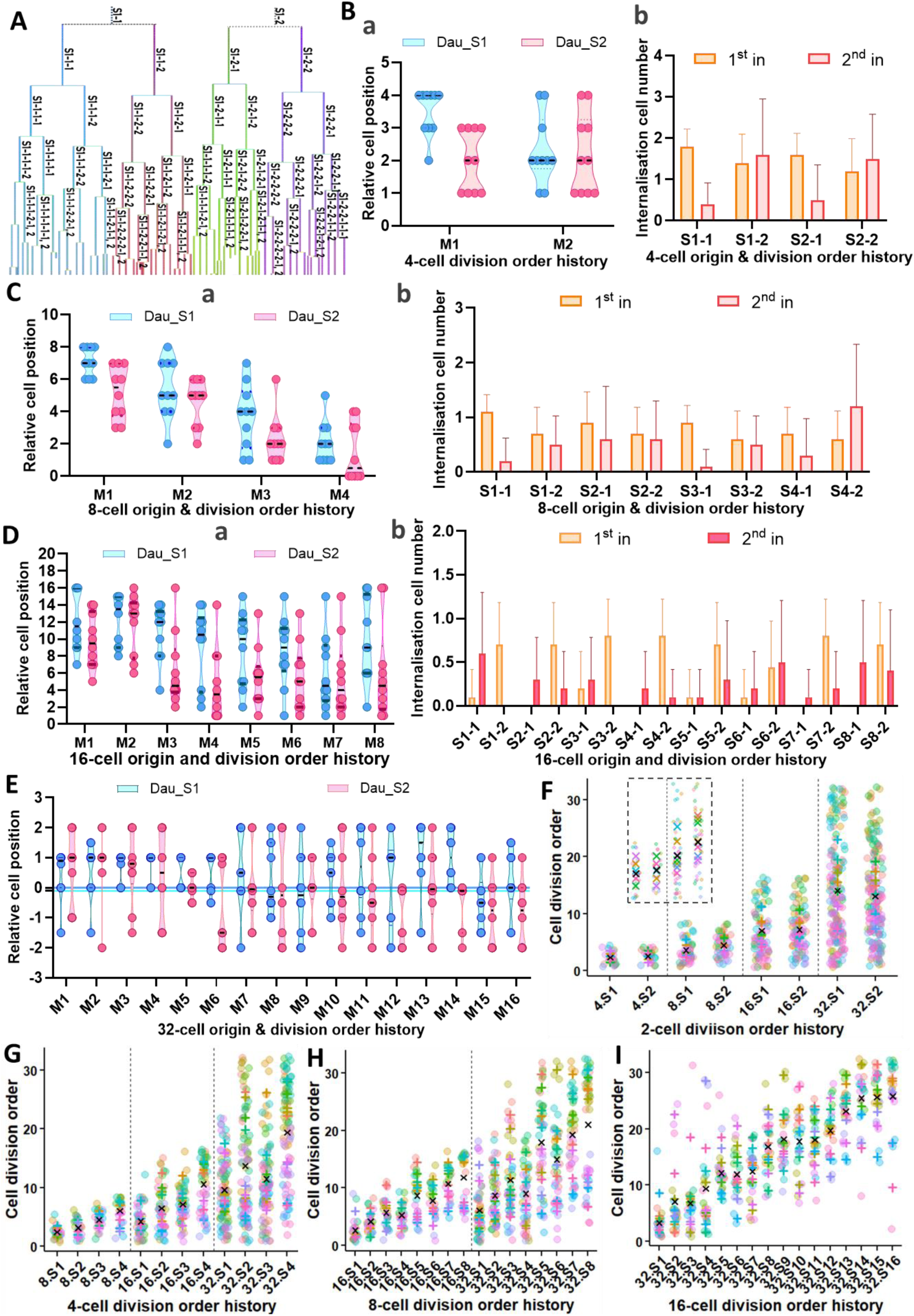
Histories and relations of cell division orders and cell positions. In all panels, “S” refer to the “sequence” (order) of cell division. **(A)** Cell tracking trees with number-and colour-coded cell division order histories. **(B-E)** Relations of cell division order histories between two consecutive generations and cell positions/internalisation for 4-, 8-, 16-and 32-cell stages. Materials and Methods show detailed approaches for position ranks in the Y-axis in (B-E). Subpanels a in (B-E): Relations of cell relative positions and order history marked by mother division order (M) from 1 (the earliest) to the latest (depending on mother cell counts) and their pairwise daughter (Dau) division orders (S1 for earlier order and S2 for the later one). Subpanels b in (B-E): Relationships between cell division order history and the 1^st^ (orange) (the 4^th^ division round) and 2^nd^ (the 5^th^ division round) cell internalisation (pink), with the first number behind S for mother division order, and the second for their daughter division order, and “cell number” on the Y-axis for cell counts. **(F-I)** Longitudinal cell division order histories without distinguishing pairwise daughters from the same mother. Each colour represents a different embryo, with the corresponding coloured cross indicating the average for each embryo. The black cross represents the overall average across all embryos. (F): Division order history of 4-cell (orange), 8-cell (yellow), 16-cell (green), and 32-cell offspring (blue) from 2-cell mothers. (G-I): Division order histories of offspring from 4-cell mothers, 8-cell mothers, and 16-cell mothers, respectively. The error bar in all panels (except for Panel A): SD. Embryo sample size: 10 in (B-E), six in (F-I). In (F-I) (from two videos across three litters). Each colour-coded line represents each embryo.

Earlier-dividing 4-cell blastomeres (S1-1) were farther from the embryo centre (position ranks: 2.3 ± 1.05 to 3.5 ± 0.7) than the later-dividing 4-cell sisters (position ranks: 2 ± 0.94 to 2.23 ± 1.29), regardless of their 2-cell mothers division orders (Figure 7Ba). Furthermore, early-dividing 4-cell blastomeres from each 2-cell mother had their descendants mainly internalised in the 4^th^ round (30 % of 60 cells from S1-1 and 27% from S2-1) compared to the slower-dividing 4-cell sisters (23% from S1-2 and 20% from S2-1), whose descendants were mainly internalised in the 5^th^ round (40% of 40 cells from S1-2 and 37.5% from S2-2) (Figure 7Bb). Daughters of earlier-dividing 2-cell mothers tended to show more notable differences in descendant numbers internalising in the 4^th^ and 5^th^ rounds than those from later-dividing 2-cell mothers (Figure 7Bb). Earlier-dividing 8-cell blastomeres from each 4-cell mother were located farther from the embryo centre (position ranks: 2.10 ± 1.20 to 7.10 ± 0.87) than their later-dividing sisters (position ranks: 1 to 5.20 ± 1.69) (Figure 7Ca). These earlier-dividing 8-cell blastomeres tended to have more descendants internalised in the 4^th^ round than in the 5^th^ round, while later-dividing 8-cell sisters tended to have more descendants internalised in the 5^th^ round, though this pattern slightly varied among all 8-cell daughter pairs. The differences in cell numbers internalised in the 4^th^ and 5^th^ rounds were more pronounced for the 8-cell descendants from 4-cell grandmothers with odd division orders (1 and 3) than those with even orders (2 and 4) (Figure 7Cb).

Early-dividing 16-cell blastomeres from each 8-cell mother were mainly positioned outside of the embryos (position ranks 6.10 ± 4.10 to 12.30 ± 2.94), while their later-dividing sisters were relatively dispersed across the embryo but locally clustered towards the inside (4.70 ± 4.30 to 11.30 ± 3.52) (Figure 7Da). Those earlier-dividing 16-cell blastomeres rarely showed completed internalisation in the 4^th^ round but had more daughters internalised in the 5^th^ round, opposite to their later-dividing pair sisters. Total numbers of internalised cells sharing ancestry with earlier-dividing 16-cell blastomeres were significantly lower (one to two cells in the 4^th^ round and two to six in the 5^th^ round) than their later-dividing pair sisters (four to eight in the 4^th^ round and zero to five cells in the 5^th^ round) (Wilcoxon matched-pairs signed rank test, *p* = 0.0156) (Figure 7Cb).

The 32-cell blastomeres from the first six dividing 16-cell mothers tended to cluster around the embryonic part, while their cousins from the other 10 16-cell mothers spread across embryonic and abembryonic parts, focusing on the ICM surface and Mu-TE (Figure 7E). Among the latter, earlier-dividing 32-cell blastomeres from each mother clustered around Mu-, P-TE, and ICM lining P-TE near Mu-TE, while their later-dividing sisters clustered from P-TE and nearby deep ICM, especially notable for descendants from the 16-cell mothers in orders 12-16 (Figure 7E). The first seven dividing 32-cell daughters almost exclusively clustered outside, cells in division orders 8-14 tended to be around the P-TE near Mu-TE and ICM surface, and the rest were mainly in the deeper ICM.

I also examined changes in division orders across generations. Compared to slower-dividing 2-cell descendants, faster-dividing 2-cell descendants were divided earlier in the 3^rd^ and 4^th^ division rounds (Figure 7F). In the 5^th^ round, both groups of descendants showed equal division orders, but in the 6^th^ round, the orders between the two groups reversed (Figure 7F). Descendants of 4-cell ancestors consistently mirrored their mother division orders in the 4^th^ and 5^th^ rounds (Figure 7G). In the 5^th^ round, this pattern persisted for the descendants of the two first-dividing and two later-dividing 4-cell blastomeres, respectively (Figure 7G). Descendants of the 8-cell blastomeres from the first two and last two division orders followed their mother division orders in both the 4^th^ and 5^th^ rounds. Descendants of mid-division ancestors reversed their orders every two orders (Figure 7H). Similar alternations were observed between 16-cell blastomeres and their 32-cell descendants (Figure 7I). The reversed historical cell division orders were in line with the changes in dynamic-to-determined cell positions at the given developmental stages.

These data show that division rates of pairwise cells decrease when they are nearer embryo centres from 2-to 32-cell stages, with earlier-dividing sisters mainly internalised earlier at 8-to 16-cell stages. Furthermore, division orders are consistently inherited from the 1^st^ to 4^th^ rounds, with minor reversals between the 5^th^ and 6^th^ rounds as cells differentiate into internal and external positions. These findings suggest that both spatial location and division historical inheritance contribute to early asymmetry events of lineage segregation.

#### 2.4.4 Impact of cell division asynchrony and synchrony on lineage allocation

Building on the above-analysed relationships between cell lineages and division orders, I explored how cell lineages at E4.5 related to division time gaps (asynchrony extent) between pairwise cells from 2-to 16-cell stages using the local smooth curve (Figure 8A,B). The asynchronous extent was measured as the absolute division timing differences of two pairwise daughter cells sharing the same mothers, sorted by division orders. In 10 embryos, TE cell counts increased from 25 to 37 as division asynchrony between 2-cell blastomere pairs changed from 20 to 60 minutes, then dropped to 27 at 100 minutes. ICM cells rose from 18 to 21 ± 3, decreased to 19 ± 1 with 20-to 60-minute asynchrony, then increased to 36 at 100 minutes. Total embryonic cell counts increased from 43 to 57 ± 2 between 20-to 60-minute asynchrony (Figure 8C). The 4-cell pair blastomere asynchrony between 20 to 60 minutes resulted in ICM counts from 9 ± 1 to 14 ± 2, then gradually declined from 14 to 10 at 180 minutes. TE cells changed around 32 cells before 60 minutes of asynchrony and then gradually decreased from 19 to 16 cells between 60 to 120 minutes, with the total cell number changing from 32 to 25 (Figure 8D). The ICM, TE and total cell counts changed around 6 ± 2, 9 ± 2 and 15 ± 2.5, respectively, during 0 to 120-minute asynchrony of 8-cell blastomere pairs, followed by a decrease within each group (Figure 8E). The ICM, TE and total cell counts changed around 3 ± 1.5, 4 ± 1.5 and 7 ± 1.5 respectively during 0 to 140-minute asynchrony of 16-cell pairs, followed by decreases despite a slight increase in TE cells (Figure 8F).

**Figure 8.**
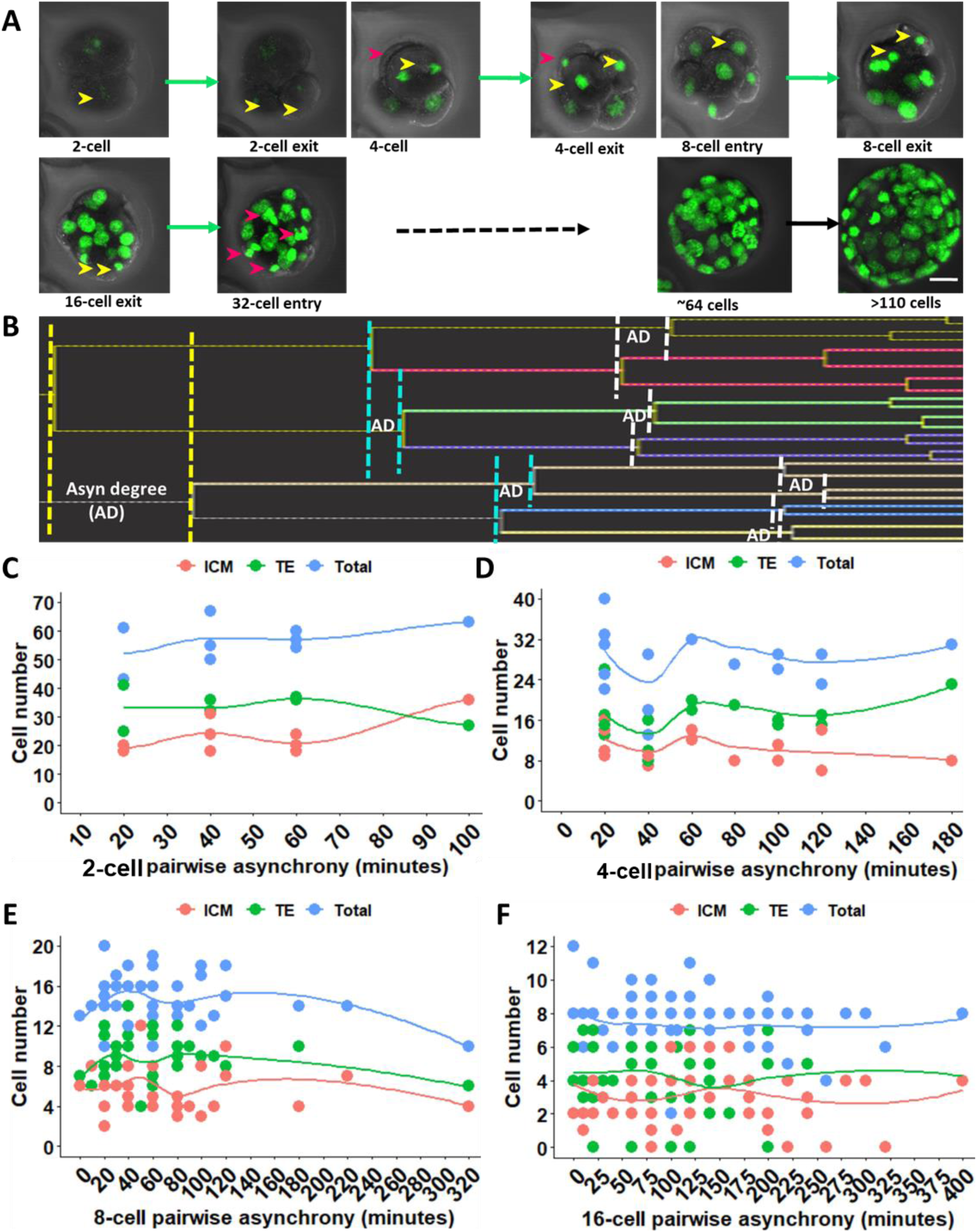
The relations between cell division asynchrony degree and cell lineage formation. **(A)** Asynchronous divisions during each division round, with yellow and pink arrows coding cells dividing at different times within the same round. The scale bar is 20 µm. **(B)** Division asynchrony degree between pairwise daughters at each division round. **(C-F)** Correlation of asynchrony degree and the counts of TE (green), ICM (pink/blue), and total embryonic cell numbers (light blue/purple) at the 1^st^ (2-cell), 2^nd^ (4-cell), 3^rd^ (8-cell), and 4^th^ (16-cell) division rounds, respectively. The plots employed a local regression (smooth curve) for fitting the data at each time point across division rounds. Embryo size: 10 embryos (from two videos across three litters); 20 to over ∼160 cells from 2-to 16-cell stages.

These data show that pairwise cells with asynchronous division differences ranging from ∼20 minutes to 100 minutes tend to contribute to relatively higher total cells, including both TE and ICM cells, during pre-and peri-implantation stages. This indicates the significant role of both asynchronous and synchronous divisions in cell lineage expansion and embryo growth.

### 2.5 Dynamics of Cell Cycle Length and Their Influence on Cell Lineage Specification

#### 2.5.1 Various cell cycle lengths within and across cell generations

One of the potential drivers of the observed cell division asynchrony extent and division orders is varied cell cycle lengths. To explore cell cycle variations, I measured cell cycle lengths of individual embryonic cells based on cell tracking trees from E1.5 to the end of long-live H2B-GFP videos (Figure 9A) (see Materials and Methods for cell cycle measurement). Embryonic cell cycle progression periods were termed the 1^st^ to 8^th^ period (refer to Appendix C for details), reflecting cell generations (Gn1-Gn8) and the 1-, 2-, 4-, 8-, 16-, 32-, 64-, and 128-cell stages, respectively (refer to Figure 5B). However, the 32-, 64-and 128-cell stages represented cell numbers for the 6^th^ and 7^th^ periods of cell cycles rather than the actual total embryonic cell number due to overlapping divisions (see Section 2.4.1 for details). Cell sample sizes are summarised in (Figure 9B).

**Figure 9.**
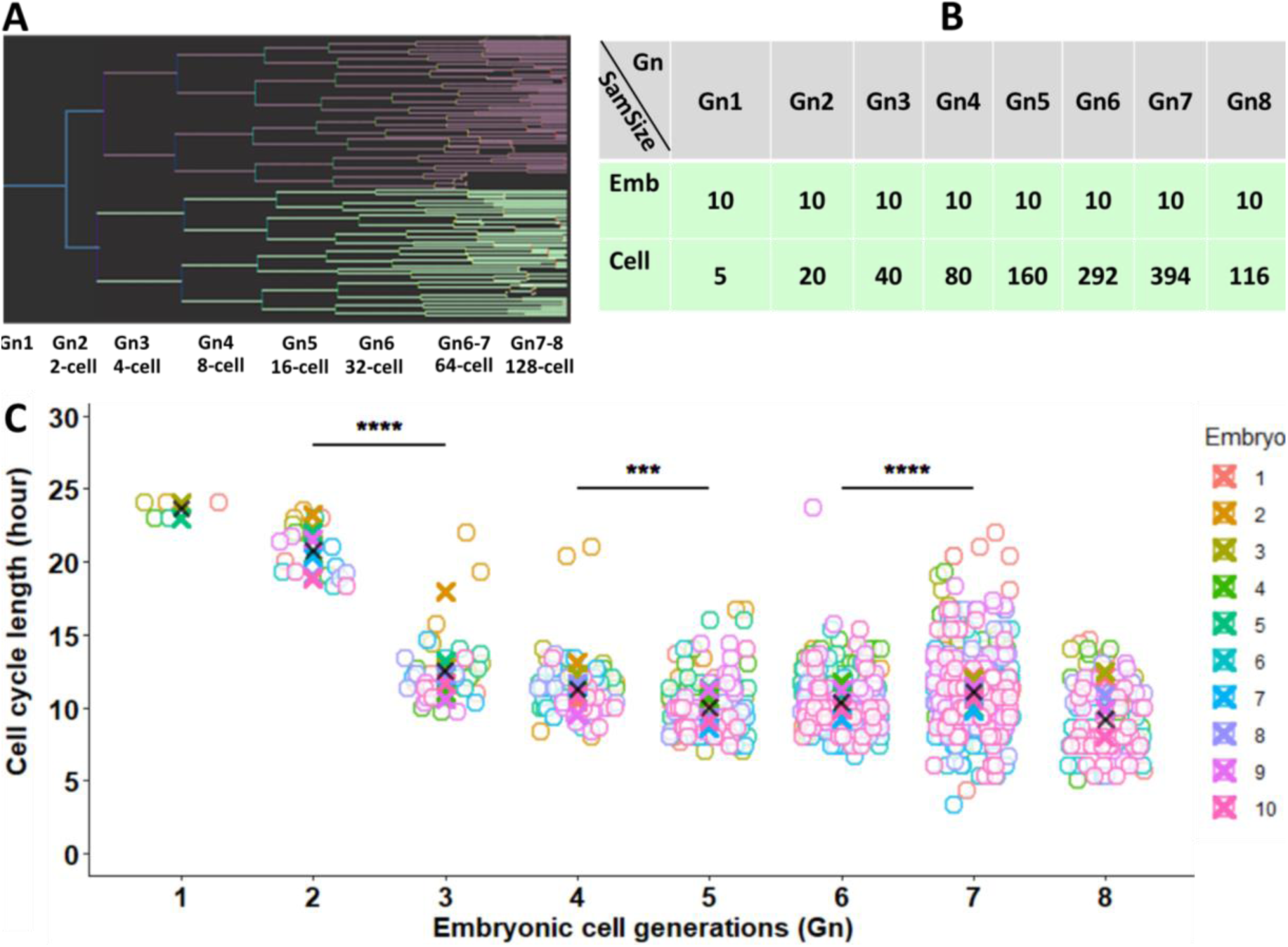
Generation definitions and cell cycle dynamics at early mouse embryo stages. **(A)** Relationships between two different systems to mark embryo developmental stages, with overlap between the start and the end of the 6^th^ to 8^th^ generations. The top level of the cell cycle tree (blue) refers to Gn1, then generations (2-8). **(B)** Analysed sample sizes of embryos and cells in each generation (Gn), with “SamSize” for sample size, and “Emb” for embryos. **(C)** Cell cycle lengths throughout the 1^st^ to 8^th^ generations and Wilcoxon matched-pairs signed rank test results with SD error bar between two consecutive generations. Each of the 10 embryos (from two videos across three litters) was colour-coded as shown in the Figure. Asterisks: \**p* < 0.05, \*\**p* < 0.01, \*\*\**p* < 0.001, and \*\*\*\**p* < 0.0001.

Initially, cell cycle lengths were 23.60 ± 0.55 and 20.80 ± 1.72 hours during the first two generations across 10 embryos, in line with previous studies (Kelly *et al*., 1978; Bischoff *et al*., 2008). The Gn3 showed a sharp decline to 12.49 ± 2.38 hours. Through the 4^th^ period, cell cycles reduced to 11.26 ± 2.01 hours, stabilising at 10.04 ± 1.99 and 10.35 ± 1.88 hours in the 5^th^ and 6^th^ periods, then increased to 11.10 ± 2.83 in the 7^th^, and decreased to 9.19 ± 2.40 in the 8^th^ periods (Figure 9C). It is worth noting a survivorship/access bias in the 8^th^ cell cycle period due to the limited video durations, meaning that only a certain subset of relatively short cell cycles was able to be observed in the 7^th^ and 8^th^ periods (Figure 9C). There were statistically significant differences in cell cycle durations between two consecutive periods except for Gn5 and Gn6 using the Wilcoxon matched-pairs signed rank test: Gn2 vs Gn3 (*p* < 0.0001), Gn4 vs Gn5 (*p* = 0.0003) as well as between Gn6 and Gn7 (*p* < 0.0001) (Figure 9C). Within each embryo, across most developmental stages, cell cycle lengths in each embryo tended to cluster into two or three distinct groups, identified by convergence regions within the plots (top and bottom, and the middle regions in the staggered plots). Notable variations in cell cycles within each generation became notable during the 4^th^-6^th^ periods, with cell cycling ranging from 8 to 21 hours, 7 to 16.67 hours, and 8 to 14.67 hours, respectively. During the 7^th^ period, the range widened from 5 to 17.67 hours, and from 5 to 15 hours in the 8^th^ period (Figure 9C).

These data illustrate continuous dynamic changes in cell cycle durations across eight generations during E1.5 and E4.5, with significant cycle differences observed at the 2-to 4-cell, 8-to 16-cell, and 32-to 64-cell stages. These dynamic changes and shifts likely reflect key regulatory events in early embryonic development, offering a foundation to explore the cell cycle relations with lineage specification and the underlying mechanisms.

#### 2.5.2 Cell cycle length variation in embryonic cells with different positions

The observed different clusters of cell cycle length distribution in each cycle period and cycle shifts across generations could be related to the unknown presence of different cell populations. To explore this potential link, cell cycles and cell positions/cell fates were evaluated (see Materials and Methods for details on cell position categorisations).

Cell cycles were compared between TE (outside) cells and ICM (inside) cells post-cavitation (Gn5-Gn8) by Wilcoxon matched-pairs signed rank test (Figure 10A). Outside cells exhibited (9.19 ± 0.78)-hour cell cycles and inside cells had (10.54 ± 1.12)-hour cell cycles in Gn5; whereas TE and ICM cells in Gn6 showed (9.976 ± 0.75)-and (10.73 ± 1.11)-hour cell cycles, respectively, with significant differences between the two groups at both stages (*p* = 0.0039 and *p* = 0.047, respectively) (Figure 10B). Cell cycles in TE and ICM groups during Gn7 and Gn8 were 10.91 ± 0.58 and 11.37 ± 1.307, 9.76 ± 2.68 and 9.53 ± 1.12 hours, respectively (Figure 10B). TE and ICM cell cycles in Gn7 exhibited an increased diversity within individual embryos. TE cell cycles varied from 7.67 to 22 hours, concentrating on shorter cycle ranges, while ICM cycles ranged from 5 to 20.33 hours, centring on the middle durations (Figure 10B). It is noteworthy that 14%-75% of cells in Gn7 and 84%-97% of cells in Gn8 were not captured for intact cell cycles. This made it impossible to definitively classify their cycle lengths as shorter or longer than 7.67 to 22 hours due to their asynchronous initiation.

**Figure 10.**
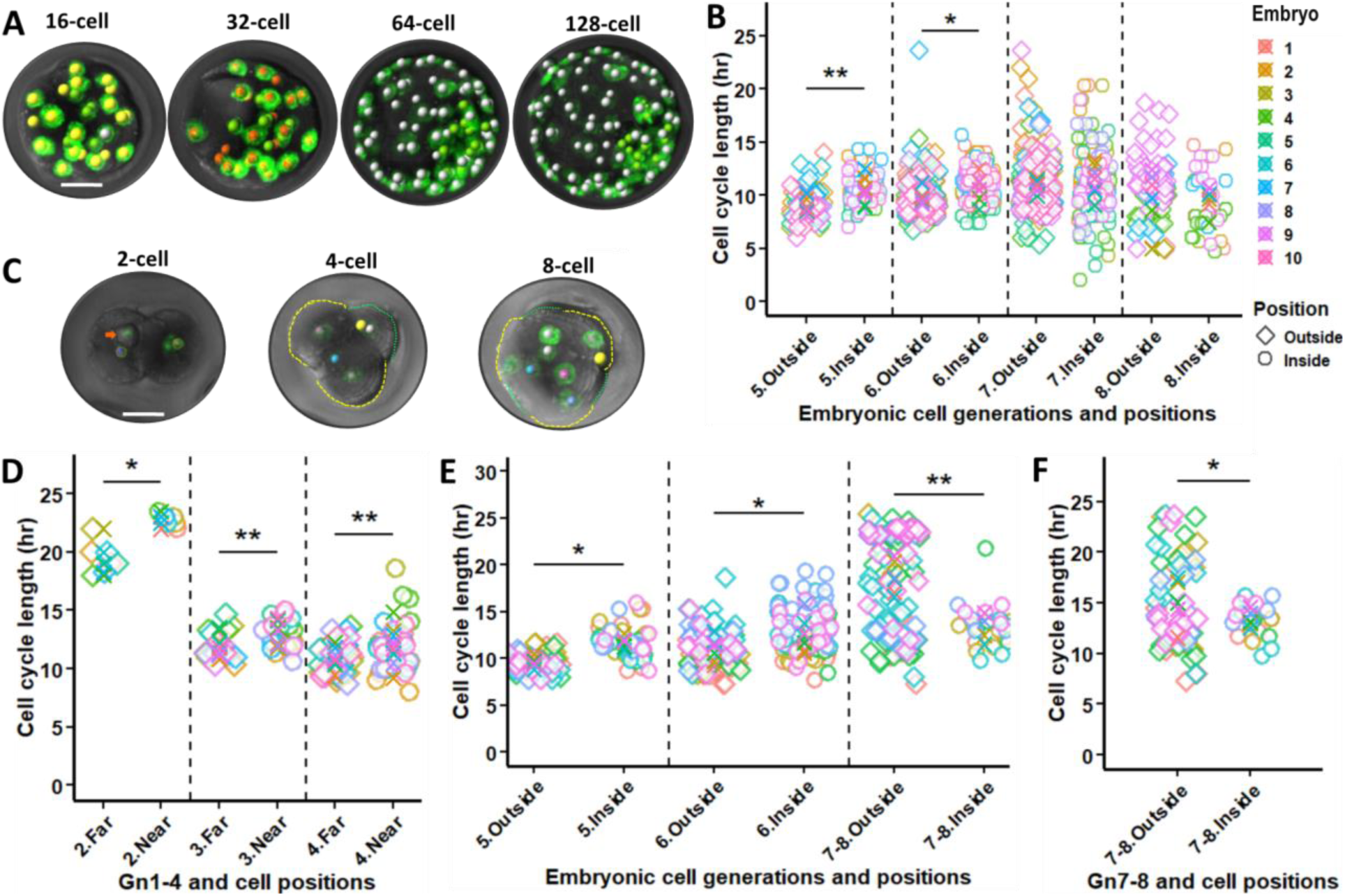
Cell cycle length and cell position. **(A,C)** Cell position categorisation. Cell positions in (Gn5-Gn8) were confirmed by cell tracking, including outside (yellow, orange and white) cells and inside cells (green). Cell positions in Gn2-Gn4 were defined by the cell nuclei distances to the nearest free-contact surface (Gn2) or to the embryonic geometrical centre (Gn3 and Gn4), with yellow for near to the embryo centre and green for far from the embryo centre. **(B,D-F)** Relationships between cell cycles and cell types featuring different cell positions, with Wilcoxon matched-pairs signed rank test results and SD error bar. The scale bar in **(A,C)** is 25 µm. Embryo sample size: 10 in (B,D), six in (E,F) (three litters). In (B-F), each of 10 embryos (from two videos across three litters) was colour-coded as shown in the Figure. Asterisks: \**p* < 0.05, \*\**p* < 0.01, \*\*\**p* < 0.001, \*\*\*\**p* < 0.0001.

Before cavitation (Gn2-Gn4), cells were categorised into near or far groups based on their nuclei distances to the nearest free-contact surface (Gn2) or to the embryonic geometrical centre (Gn3 and Gn4) (Figure 10C). In Gn2, near cells displayed significantly longer cell cycles by 3 hours than far cells (*p* = 0.0156) (Figure 10D). In Gn3, cell cycles were 11.88 ± 0.93 hours for the far group and 12.87 ± 0.97 hours for the near group (*p* = 0.0059) (Figure 10D). In Gn4, cell cycles were 10.80 ± 0.77 and 11.92 ± 0.55 hours in the far and near groups, respectively (*p* = 0.0098) (Figure 10D). Next, I investigated the potential subcellular factors affecting cell cycles. I found that fast-dividing 2-and 4-cell blastomeres had consistently more nucleoli than their sister or cousin cells at early cell cycle stages: 4 ± 1 vs 5 ± 1 for 2-cell blastomeres, 2.7 ± 0.58, 1.7 ± 0.58, 2.7 ± 0.58, and 1.3 ± 0.58 for 4-cell pairwise blastomeres.

Similar analyses were conducted on embryos in short videos. In Gn4, cells in the far group had shorter cell cycles by 1.5 hours than the far group. In Gn5 and Gn6, cell cycles were significantly shorter in outside/TE cells than those in inside/ICM cells (*p* = 0.031 and *p* = 0.031), similar to the results from long videos (Figure 10E). For cells potentially overlapping between Gn7 to Gn8, cell cycles were significantly different between TE (18.68 ± 1.75 hours) and ICM cells (13.37 ± 1.40 hours) (*p* = 0.002) (Figure 10E). However, unlike the comparison in Gn7 from the long videos, cell cycle lengths switched between these two cell populations at Gn7-Gn8 in short videos, with the TE group having longer cell cycles and ICM groups having shorter cell cycles (Figure 10E). Comparing cell cycle lengths at each stage between long and short H2B-GFP embryo videos showed little to no differences. Cell numbers in the short videos were then normalised based on cell numbers in the long videos, excluding data from cells that had progressed beyond the developmental stages represented by total cell numbers of embryos in the long videos. The normalised data analysis showed that TE cells had longer cell cycles (15.03 ± 4.23 hours) than ICM cells (13.15 ± 1.62 hours) (*p* = 0.042), with TE cell cycles ranging from 7 to 23.33 hours and ICM cell cycles ranging from 9.25 to 16 hours (Figure 10F).

These data highlight that cells far from the embryo centre have shorter cell cycles than cells near the embryo centre during Gn2 to Gn6, with shifts in Gn7 to Gn8, underscoring that cell positions dynamically influence embryonic cell cycles as embryos grow over time.

#### 2.5.3 Cell cycles in different subpopulations of outside and inside cells

Given the above-mentioned diverse cell cycles in later cycle periods, it was crucial to investigate the topography of these cells within the ICM and TE compartments. I produced a heatmap for the Gn6 duration, chosen for its comprehensive cell cycle tracking and a higher likelihood of ICM and TE sublineage formation (Figure 11A). The heatmap suggested that TE cell cycle lengths were linked to their position relative to the embryonic-abembryonic axis (Figure 11A). These TE cells were then categorised into Mu-TE and P-TE cells in Gn5 to Gn8 (Figure 4.11, Panel B), with cell sample size summarised in (Figure 11H). Cell cycles of P-TE and Mu-TE cells in Gn5 were 10.03 ± 1.20 and 8.98 ± 0.87 hours, respectively. In Gn6, cell cycles of P-TE, and Mu-TE cells were 10.26 ± 0.94 and 9.18 ± 0.61 hours, respectively. However, in Gn7, P-TE and Mu-TE cells had cell cycles of 10.77 ± 1.06, and 24.63 ± 5.23 hours, respectively. These comparisons showed statistically significant differences (Wilcoxon matched-pairs signed rank test, *p* = 0.0039, *p* = 0.0039, and *p* = 0.0117 across three paired groups, respectively) (Figure 11C). It is noteworthy that shifts of cell cycle lengths between P-TE and Mu-TE across Gn5-Gn7 occurred as a gradual cline through spatially intermediate TE cells between well-located P-TE and Mu-TE. Cell cycles of P-TE and Mu-TE in Gn8 showed a similar trend to those in Gn7 (Figure 11C). Remarkably, within Mu-TE cells in Gn6, Gn7 and Gn8, cells with shorter cell cycles were surrounded by loosely packed neighbouring cells while their pairwise sisters with longer cell cycles were surrounded by more crowded neighbouring cell environments.

**Figure 11.**
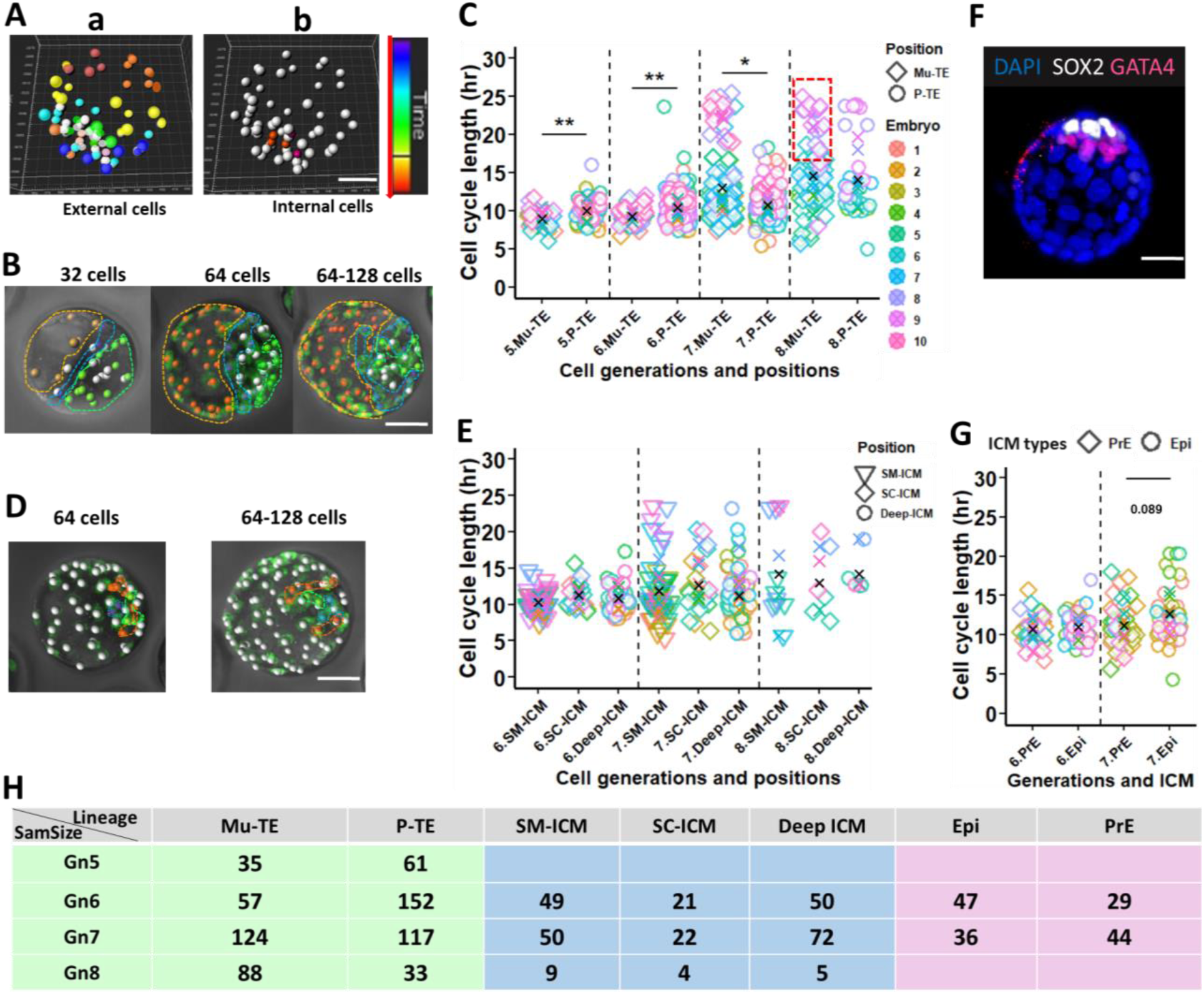
Cell cycle length in subpopulations of TE and ICM. **(A)** The heatmap of the track duration of outside cells in IMARIS, with red spots for cells with longer track duration, and blue to red spots representing the track duration of cells gradually increases. **(B)** TE cell categorisation, including Mu-TE (orange), and P-TE (green and blue). **(C)** TE cell cycles. **(D)** ICM cell categorisation based on cell tracking, including cells at the margin of the ICM surface (SM-ICM, red), cells at the centre of the ICM surface (SC-ICM, green) and deep ICM (blue). **(E)** ICM cell cycles. **(F)** ICM cell categorisation based on immunostained markers with Hoechst-stained nuclei, including Epi (Sox2-white) and PrE (Gata4-red) cells. **(G)** Epi and PrE cell cycles. In (C,E,G), each colour refers to each embryo, and the Wilcoxon matched-pairs signed rank test was applied to test cell cycle differences between different cell types in (C,G). The Scale bar in (A,B,D,F): 30 µm. The error bars in (C,E,F): SD. **(H)** Analysed sample sizes of embryonic cells. In (C,E,G), each of the 10 embryos (from two videos across three litters) was colour-coded as shown in the Figure. Asterisks: \**p* < 0.05 and \*\**p* < 0.01.

I also produced a heatmap showing ICM generation durations (Figure 11Ab), which indicated a link between cell cycle lengths and cell positions at the deep ICM, and at the margin or centre of the ICM surface. Thus, ICM positions were categorised into deep, surface margin (SM), and surface centre (SC) groups (Figure 11D,E). In Gn6 and Gn7, the deep ICM, SM and SC groups displayed cell cycles between 10.12 ± 0.84 and 11.18 ± 0.96 hours, 12.25 ± 2.89 and 13.47 ± 2.64 hours, respectively (Figure 11E). In Gn8, the deep, SM and SC ICM groups exhibited cell cycles of 15.04 ± 5.60, 11.25 ± 5.54 and 13.62 ± 6.207 hours, respectively (Figure 11E). Based on cell cycle tracking and cell matches between two sets of images including live-embryo videos and their corresponding immunostained images (Figure 11F), I analysed the latest cell cycle of Gata4-positive and Sox2-positive cells at the end of long 4D live-embryo videos. While cycle lengths did not significantly differ between Gata4-positive and Sox2-positive cells, their distributions varied (Figure 11G). In Gn6, cell cycles of Sox2 cells were similar, while in Gn7, cells cycles of Sox2 cells (12.57 ± 2.00 hours) tended to be longer than those of Gata4 cells (11.47 ± 1.71 hours) (*p* = 0.089) (Figure 11G).

These data highlight that Mu-TE cells have shorter cycles than P-TE cells in Gn5 and Gn6, but this trend reverses in Gn7 and Gn8, particularly due to lengthened Mu-TE cell cycles. Similarly, while PrE and Epi precursor cells have comparable cycles in Gn6, PrE cycles tend to shorten relative to Epi cycles in Gn7. Furthermore, loosely packed embryonic cells in each population possess short cell cycles, particularly notable among pairwise sister or cousin cells. These findings underscore cell cycle shifts likely correspond to transient lineage separation, neighbouring cell packing, and sustained maintenance of cell lineages over time.

#### 2.5.4 Cell cycle length histories and their links with cell lineages

To understand how cell cycles correlated with various cell populations, I examined cell cycle histories across different cell generations and their relations to cell positions (Figure 12A). Initially, in Gn3, the descendants of the 2-cell blastomere with short cycles switched to longer cycles, while those from long-cycle 2-cell ancestors changed to relatively short cycles. In Gn4, the cell cycle trend of 8-cell descendants mirrored the 2-cell ancestors. In Gn5, the trend switched to be opposite of that in Gn2 (Figure 12B). In Gn6 and Gn7, the general cell cycles of descendants from both 2-cell blastomeres were similar. However, in 70% of individual embryos, cell cycles in Gn6 reverted to the pattern observed in 2-cell blastomeres. In contrast, cell cycles in Gn7 shifted in the opposite pattern, a trend that became more remarkable in Gn8 (Figure 12B).

**Figure 12.**
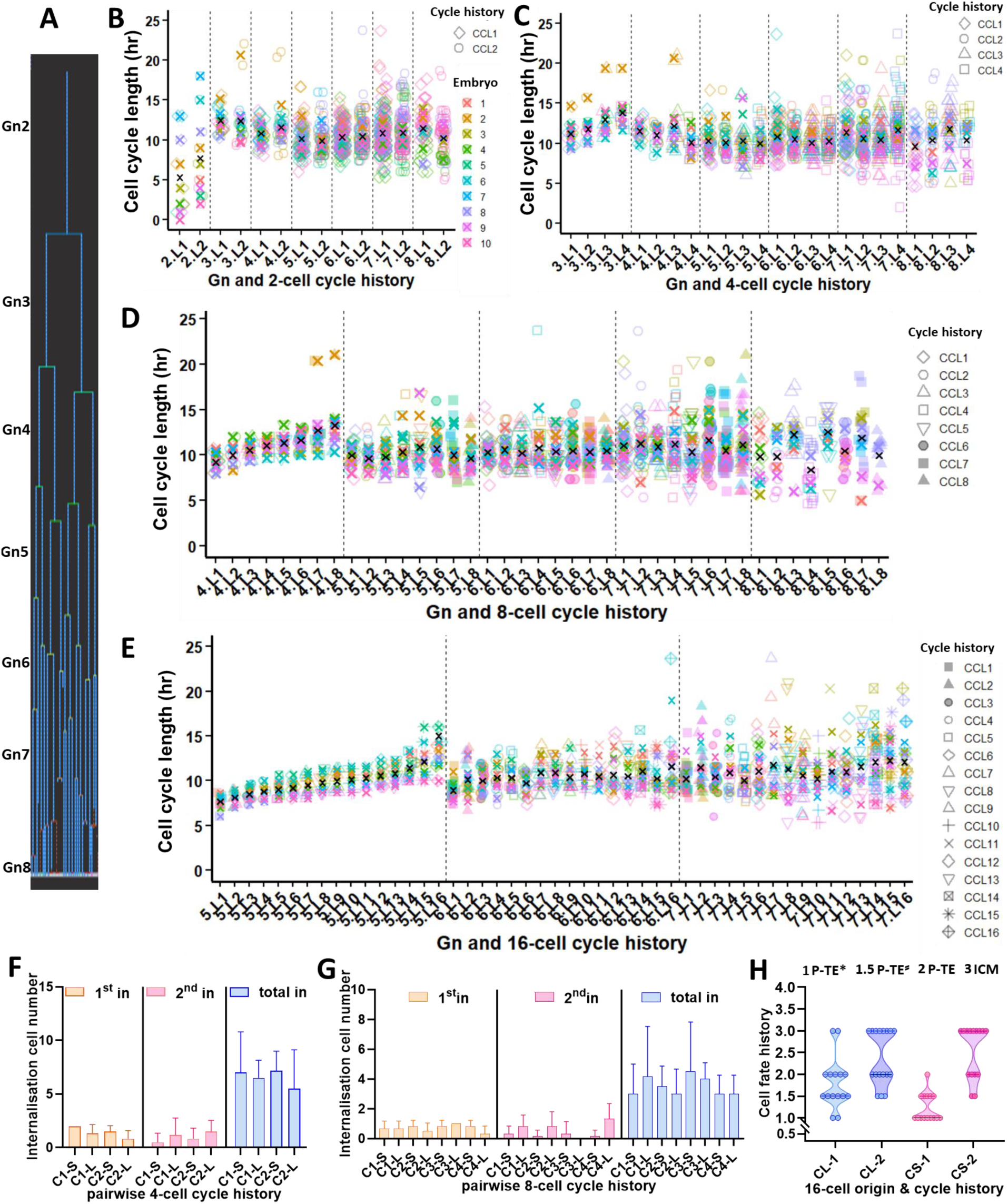
Cell cycle histories within embryos over early developmental stages. **(A)** Representative of cell generations and cell cycle histories. In **(B-E)**, each colour codes each embryo. **(B)** Cell cycle histories of descendants from short-(L1) and long-cycle (L2) 2-cell ancestors. **(C-E)** As Panel B but for descendants from 4-cell, 8-cell and 16-cell ancestors arranged from the shortest to longest cell cycle length categories (CCL). **(F)** Internalised descendants in the 4^th^ (1^st^ in) and 5^th^ (2^nd^ in) division rounds and total internalised cells (blue) from pairwise short-and long-cycle 4-cell ancestors. **(G)** As Panel F but form 8-cell ancestors. **(H)** Cell fate allocations from short-and long-cycle 16-cell external ancestors, including P-TE* (those near Mu-TE), P-TE⸗ (located between P-TE at the embryonic pole and P-TE near Mu-TE), P-TE at the embryonic pole and ICM (3), with SD error bar. CL indicates a long cycle. CS denotes a short cycle, with “1” and “2” for short-and long-cycle pairwise daughter IDs. The error bars in (B-H): SD. Embryo sample sizes: 10 (B-F), and six (F-H). In (B-E), each of 10 embryos (from two videos across three litters) was colour-coded as shown in the Figure.

The 4-cell ancestors were arranged from the shortest to longest cell cycles. In Gn4 and Gn5, descendants of the odd-category 4-cell ancestors (1 and 3) inherited their ancestor cell cycle trend, but descendants of the even-category 4-cell ancestors showed shorter cell cycles than their cousins. In Gn6, the trend shifted to be opposite to the pattern observed in 4-cell ancestors (Figure 12C). In Gn7, descendants of the last two-category 4-cell ancestors mirrored the initial orders of the 4-ancestor cycle. In Gn8, the first three-category cycles reverted to the 4-ancestor cycle pattern (Figure 12C). The Gn5 descendants of the first and the second four-category cycled 8-ancestors showed a descending trend of cell cycles (Figure 12D). In Gn6 and Gn7, the descendants of 8-cell ancestors with odd-and even-numbered cycle categories had slightly shorter and longer cell cycles, respectively, opposite to the pattern in Gn8 (Figure 12D). In Gn6, the descendants from 16-cell blastomeres had shorter cell cycles in odd categories and longer cell cycles in even categories, opposite to the pattern observed in Gn7 (Figure 12E).

I observed distinct cell cycle patterns among four cousins sharing grandmothers in Gn2 to Gn6. In each generation, one sister pair consistently showed more notable cell cycle differences than their cousins (the other sister pair), with the first pair having the longest and shortest cycles and the other pair showing medium cycles. Interestingly, each of four cousin cells cohorts exhibited approximately 1:2:1 ratio for the longest, medium and shortest cycles, suggesting a possible binomial segregation and inheritance of a cell cycle-determining factor across two sequential generations. These differences in cell cycles between pairs of sisters and their cousins appeared to be influenced by their position allocation within embryonic local structures. To confirm this observation, I analysed cell cycle histories of pairwise daughters in each generation (categorised by their mother cycle lengths), and their descendent internalisation and lineage allocation by the late 32-cell. The short-cycle 2-cell blastomeres formed 52% of 157 total ICM while their longer-cycle sisters formed 48% of ICM cells. In Gn3, the descendants of each short-cycle 4-cell blastomere tended to internalise more in the 4^th^ division round, forming 54.15% of 157 ICM; whereas the descendants of their long-cycle sisters appeared to internalise more in the 5^th^ round, forming 45.85% of ICM cells (Figure 12F). The descendants of short-cycle 8-cell blastomeres tended to internalise slightly more in the 4^th^ round (56% of the 34 1^st^ internalised cells) than their respective long-cycle sister pairs (44%), which appeared to internalise more in the 5^th^ round (75% of the 24 2^nd^ internalised cells) (Figure 12G). No clear differences or trends were observed in total ICM cells from each pair of 8-cell daughters (Figure 12G).

The 16-cell blastomeres that had not yet differentiated were classified into long-and short-cycle cells between pairwise sisters or 4-cell cousins across six embryos. Of the descendants from 15 long-cycle 16-cell blastomeres, 74% of short-cycle daughters formed P-TE except those near Mu-TE cells, with the rest forming ICM and P-TE cells near Mu-TE cells, while all long-cycle daughters formed ICM and TE cells near the embryonic part (Figure 12H). Of the descendants from 15 short-cycle 16-cell blastomeres, 93.3% of short-cycle daughters formed P-TE cells except those located in the embryonic pole, while 60% of long-cycle daughters formed ICM cells and 40% formed the embryonic part (Figure 12H). The short-cycle 32-cell daughters from short-cycle 16-cell blastomeres formed more P-TE cells near Mu-TE than those from their long-cycle 16-cell sisters or cousins, which formed more embryonic pole cells (Figure 12H).

These data show that cell cycle length is passed on either consistently or reversely across embryonic generations, depending on odd-or even-numbered cycle categories and embryo stages. My results show that cells with fewer surrounding neighbours have shorter cycles. The ancestors or descendants of these cells are linked to earlier internalisation and a propensity for ICM formation. These findings underscore the influence of cell position on shaping non-linear cell cycle histories within both the overall embryonic cell population and individual sub-lineages over time.

### 2.6 Dynamic Tango between Cell Division Orders and Cell Cycle Lengths

Following the analysed cyclic dynamics in cell cycle histories, I examined cell cycle changes with division order. In Gn3, cell cycles increased linearly with cell division orders (Figure 13A). In Gn4, they increased for the first five division orders but declined from orders 6 to 8 (Figure 13B). Cell cycles in Gn5 and Gn6 showed temporary peaks every three orders, with cycles continually increasing at higher orders (14 to 16 in Gn5, 25 to 32 in Gn6) (Figure 13C,D). In Gn7, cell cycles generally increased with division orders, peaking every two to three orders before order 24, every six orders before order 47, and then every two to three orders again (Figure 13E).

**Figure 13.**
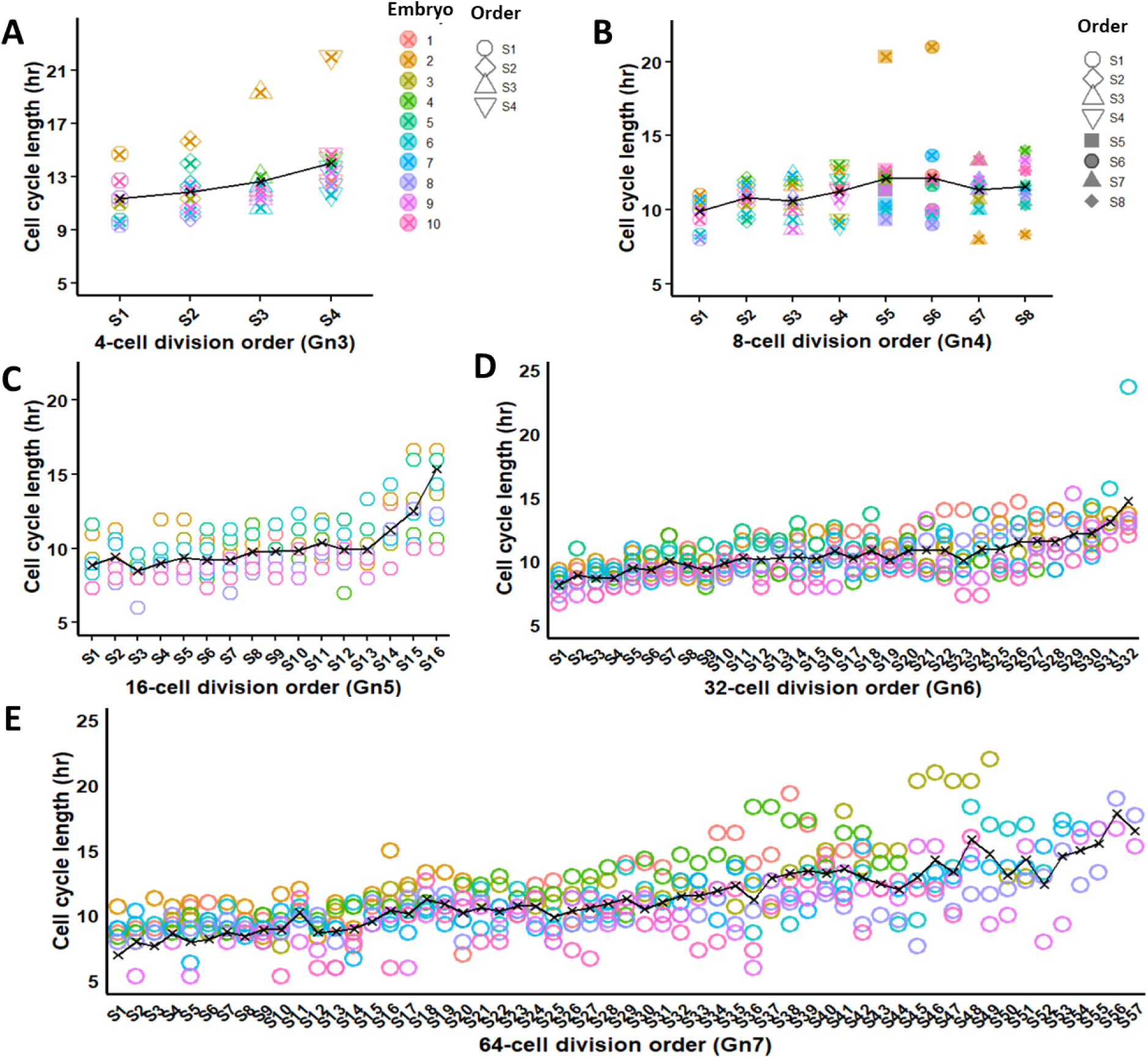
Relationships between division orders, cell cycle, and cell lineages. **(A-E)** Cell cycle changes and Kruskal-Wallis test results with SD error bar based on cell division orders (S) in Gn3 to Gn7. Cell division sequences (orders) were colour-coded as shown in each panel. Embryo sample size: 10 (from two videos across three litters). Each of the 10 embryos was colour-coded as shown in the Figure.

These data reveal a linear increase in cell cycle durations with cell division orders in Gn3, and dynamic non-linear relationships in Gn4 to Gn7-characterised by alternating patterns and periodic peaks of cycles with division orders. This indicates a structured yet dynamic interplay of cell cycle durations and division orders.

### 2.7 Interplay of Cell Positions, Cell Divisions, Cell Cycles and Cell Lineages

Overall, descendants of early-dividing (short-cycled) 2-cell blastomeres (positioned relatively far) internalised more in the 4^th^ round, forming higher numbers of ICM cells by the 32-cell stage. This trend continued through the 4-cell, 8-cell, and up to 32-cell stages (Figure 14A-C). Within each pair of consecutive generations, the descendants of early-dividing short-cycle 4-cell blastomeres (positioned relatively far) internalised more in the 4^th^ round, as did their pairwise sisters but in the 5^th^ round. This pairwise trend persisted over 8-to 16-and 16-to 32-cell stages (Figure 14A-C). The early-dividing 8-and 16-cell ancestors produced short-cycle Mu-or P-TE cells around the initiated cavity, paired with other long-cycle TE cell sisters, or short-cycle TE cells paired with long-cycle ICM cells internalised in the 4^th^ division round. The late-dividing 8-and 16-cell blastomeres generated short-and long-cycle P-TE cells, or short-cycle TE paired with long-cycle ICM cells internalised in the 5^th^ division round. At the early 32-cell stage, early-dividing cells continued to produce short-cycle P-TE cells near Mu-TE and short-cycle margin ICM cells, while their late-dividing counterparts produced long-cycle TE and ICM cells (Figure 14A-C). At the late 32-cell stage, the early-dividing cells generated short-to medium-cycle P-TE and deep ICM cells, with long-cycle pairwise sisters producing Mu-TE/P-TE (near Mu-TE) and ICM/P-ICM (near Mu-TE) cells (Figure 14A-C). Such patterns of cell origin, cell cycle, cell position and cell lineages were also applied to 64-cell stages (Figure 14A-C). Furthermore, Mu-TE cells enclosed in a crowded neighbouring environment had shorter cell cycles than their pairwise sisters originating from the same mother cells (Figure 14Ab, 32-64 cells).

**Figure 14.**
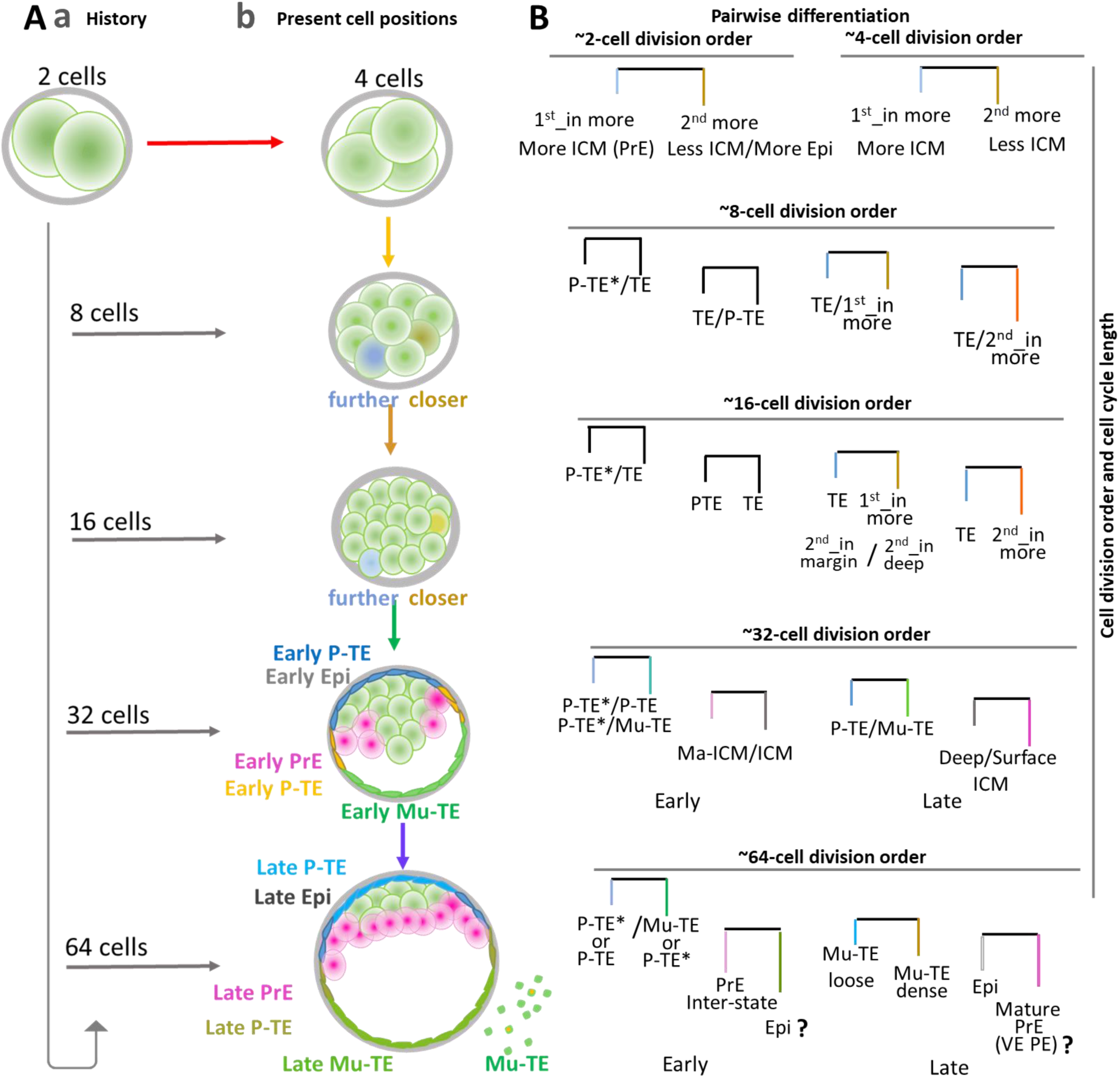
Schematic of Relationships between division orders, cell cycle, and cell lineages. **(Aa,b)**: History of embryonic cell origins and differentiation as well as cell positions and lineages, with outside cell (blue), cells near the embryo centre (dark yellow) by morula, Epi (green), PrE (pink), P-TE (blue), P-TE near Mu-TE (orange), and Mu-TE (green) at 16-and 32-cell stages. At 32-64 stages, yellow marked green Mu-TE cells represent pairwise Mu-TE cells, with the remaining green cells indicating neighbouring cells. **(B)** Relations between cell division order, cell cycles, cell position-marked cell lineages, with the same colour-coding as shown in (Ab), and question mark denoting the deducted conclusion based on the study results.

### 2.8 Cell Cycle and Morphological Events in Mouse Embryos

Drawing on the observed dynamics of cell cycle lengths and their relations with cell positions, populations/lineages and their historical inheritance, I examined cell cycle phases and their effects on embryonic cell activities. Due to resource restrictions, a detailed time-lapse examination of cell cycle phases was unfeasible in this study. As a solution, I explored the potential relations between approximate phases of cell cycles and different stages of embryo compaction, initiation of cavitation and embryo hatching. Across five embryos, compaction initiation occurred early during the 4-cell to 8-cell transition, approximately 60 ± 40 minutes post-cell divisions, and completed at the cell cycle phases of 6.5 ± 1.65 hours post-cell divisions. Embryonic cells retained such closest attachment for around 2.17 ± 1.66 hours before they exited from the 8-cell stage. The compaction was, therefore, initiated at the early phases of the cell cycle, *i.e.*, circa the one-fifth stage of total cell cycle durations. Cells near early cavities began their cycles ∼0 to 2.3 hours post the start of the 5^th^ division rounds, with early cavity expansion occurring when these cells progressed at the one-fourth stage of their cycles (∼6.7 to 10 hours). Embryonic hatching initiation often coincided with early cell cycle phases, occurring 20 minutes to one hour after entering a new cycle, which constituted approximately one-fifth of the cycle length (7.0 ± 1.7 hours).

These data show that cells enter their early cell cycle phases when initiating the critical morphological events during pre-and peri-implantation stages, highlighting the coordination between cell cycle progression and transitions of developmental events.

## 3 Discussion

This work examined the spatial and timing frameworks coordinating embryo development during pre-and early peri-implantation stages. The results revealed that dynamic historical and present spatial relationships among blastomeres (2-, 4-, 8-and 16-cell as well as later stages) directed asymmetric cell lineage development. Asynchronous cell division orders across cell generations, together with cell positions, influenced cell lineage trajectories. Moreover, cell cycle durations are influenced by cell positions and linked with cell fate decisions through interaction with cell spatial cues. Notably, these developmental cues emerged more prominently between pairwise sister cells across generations. The dynamic intricate orchestration between cell positions, division orders, cell cycles and their histories all together regulated cell lineage specification and inter-lineage interactions over time.

### 3.1 Historical Spatial Cues of Embryonic Cell Origin, Differentiation and Morphogenesis

My study demonstrated that early blastomeres produced descendants that formed different cell populations with diverse cell fates based on cell position cues, while still exhibiting relatively common temporal and spatial features and forming local cooperative cell clusters. The clustering community of cells, differentiated trajectories of cell fates, and cell heterogeneities within and across cell communities are coordinated by spatial, historical properties of embryonic cells, as discussed below.

Cell lineage specification history in this study showed that ICM and TE cells exhibited unequal contributions from each 2-and 4-cell blastomeres. These differences were associated with their relative positions to the embryo centre. Specifically, 2-cell blastomeres relatively farther from the embryo centre form more ICM cells at E3.75, with fewer TE but more PrE descendants at E4.5, suggesting position-associated asymmetrical developmental trajectories of 2-and 4-cell blastomeres. Furthermore, P-TE, Mu-TE, Epi, PrE and their potential subpopulations did not fully equally stem from each 2-and 4-cell blastomere. These findings suggest that 2-and 4-cell blastomeres already possess distinct differential potentials for various cell lineages, driven by blastomere positions. My findings enrich our understanding of spatial historical origins of different cell populations, extending previous studies on the first cell lineage (ICM and TE) to their sublineages (Piotrowska-Nitsche *et al*., 2005; Soszyńska *et al*., 2019; Yao *et al*., 2019), aligning with the finding in human embryo (Junyent *et al*., 2024). My work from non-invasive data provides reliable supports for distinct development of 2-and 4-cell blastomeres reported by Krawczyk *et al*. (2021) where embryonic cells were dissociated. My study also displayed a related corresponding pattern between PrE and TE cells from the same 2-and 4-cell blastomeres, further indicating that PrE and TE cells may originate from the same precursors at 2-and 4-cell stage, supporting a previous work showing that TE-inhibited ICM cells could form more Epi and less PrE cells (Mihajlović *et al*., 2015).

My findings on the spatial history of morula stages suggest asymmetric development of 8-and 16-cell blastomeres in cell internalisation, cavity initiation, TE/ICM and Epi/PrE specifications, depending on blastomere dynamic positions and their histories. These spatial cues may be reflected in cell polarity during cell fate decisions, concordant with a previous study by Niwayama *et al*. (2019), who reported competitive relationships between cell shape, cell polarity and cell fates. My work demonstrated the asymmetry of the internal and external spatial allocation during 8-to 16-cell and 16-to 32-cell transitions, and the relative symmetry of total external and internal cell occupation at the beginning of 32-cell stages. These findings expand the previous findings on cell internalisation (Anani *et al*., 2014; Maître *et al*., 2016) by quantitatively detailing dynamic histories of cell internalisation and externalisation, enhancing our understanding of the influence of spatial histories at the morula stages on cell lineage specification and morphogenesis. After the 32-cell stage, migrating pioneer PrE cells and hatching breaker TE cells shared common progenitors traced back to the 8-cell or earlier stages, suggesting the historical origin between certain TE and PrE cells and their interlineage interaction.

Overall, this study reveals that blastomeres integrate historical spatial cues to drive asymmetric cell allocation, developmental trajectories, lineage origin and differentiation.

### 3.2 Position-Driven Cell Division Orders Influence Cell Lineage Specification

This work demonstrated interesting specific 24-hour time patterns of embryonic cell divisions across generations. These findings provided the clear and consistent references of embryonic cell numbers and with informed 24-hour timings of cell divisions, beneficial for time management of experiments on mouse embryo collection based on different study aims.

My findings indicated a specific history of cell division orders across Gn2 (2-cell stage) to Gn6 (32-cell stage), influencing cell lineage specification. Early-dividing 2-, 4-, and 8-cell blastomeres, often on the embryo periphery, tended to internalise their daughters more in the 4^th^ division round and produced more ICM cells than their late-dividing sisters that located relatively inside and internalised descendants more in the 5^th^ round. This principle was more dominant among pairwise sister cells. Furthermore, early dividing 16-cell blastomeres had pairwise sisters internalised in the 4^th^ division, while late dividing 16-cell cousins had their daughters internalised in the 5^th^ division. These results explain and provide insights into previous findings on earlier-dividing 2-cell blastomeres forming dominant embryonic part (Piotrowska *et al*., 2001), biases of 2-and 4-cell blastomeres towards ICM and TE (Tarkowski and Wróblewska, 1967; Krawczyk *et al*., 2021), and cell position-polarity roles in TE/ICM differentiation (reviewed in Yang, 2025a). My study also showed that cells from early-dividing 2-and 4-cell ancestors tended to form more ICM with more PrE cells while late-dividing 2-and 4-cell ancestors formed more Epi cells. While 8-and 16-cell ancestors in middle division orders contributed relatively more to Epi cells; PrE origins alternatively varied in division orders between pairwise daughters but earlier-dividing sisters formed Mu-TE (near P-TE) and marginal ICM cells (indicative of potential PrE cells). The 32-cell blastomeres maintained similar patterns of cell division orders and cell differentiation. This suggests that early-internalised cells in both internalisation rounds predominantly tend to form more PrE, extending a study by Morris *et al*. (2010) on ICM cell internalisation and its relations to Epi/PrE differentiation. The current work also showed that while both earlier and later-dividing cells in the 4^th^ division round could produce protrusion-featuring descendants, early-internalised cells at 16-and 32-cell stages tended to exclusively form PrE cells with protrusions, indicating the non-identical precursor presence of PrE subpopulations (PE) as early as the 8-and 16-cell stages. Grabarek *et al*. (2012) reported that higher plasticity of differentiation of PrE cells than Epi cells, supporting the notion in this study that PrE and certain PE cells from earlier-dividing blastomeres in each round may be less differentiated (at least in certain properties) than their later-dividing cousin cells at the same stages.

### 3.3 Asynchronous Division Roles in Lineage Specification

My research revealed that asynchronous division was the predominant pattern of embryonic cell divisions throughout the pre-and early peri-implantation, corroborating the findings reported by Kelly *et al*. (1978). Here, I demonstrated an occurrence of continuous asynchronous division without obvious non-division internal across generations around the 40-to 50-cell stage. Cell division synchrony gradually increased as embryos grew. My result showed that asynchronous division degree between pairwise daughters at 20-60 minutes in 2-and 8-cell stages, and 20-100 minutes at 4-and 16-cell stages tend to result in relatively more ICM and TE and total embryonic cell numbers. These results indicate a critical role of both division asynchrony and synchrony in regulating cell fate and lineage expansion. My findings extend a study by Mashiko *et al*. (2022), who stated that asynchrony of cell divisions comprised ICM formation and led to embryo abortion in mouse and human embryos. My work also supports and extends previous findings (Cruz *et al*., 2012), showing that asynchronous divisions could aid in assessing embryo viability, with specific thresholds for asynchrony identified as 1.0 ± 0.5 hours for the 2^nd^ generation and 1.1 ± 0.5 hours for the 3^rd^ generation. Overall, my study sheds light on the threshold of asynchronous degree as a parameter for assisting in the evaluation of cell fate decisions and cell lineage during pre-and early peri-implantation embryo development.

### 3.4 Cell Cycle Duration Dynamics in Early Mouse Embryos

This study longitudinally demonstrated embryonic cell cycle lengths from 2-cell to peri-implantation (> 128-cell) stages, addressing the gap of previous studies on cell cycle lengths up to the 32-cell stage (Kelly *et al*., 1978; Bischoff *et al*., 2008). Kelly *et al*. (1978) and Bischoff *et al*. (2008) reported the first two cell cycle lengths to be ∼22-24 hours, while the following cell cycle lengths to be ∼8-10 hours by the 32-cell stage. My study observed wider cell cycle ranges during the 3^rd^, 4^th^ and 5^th^ generations (with the maximum cycle of 14, 16.67, and 14.67 hours, respectively). Kelly *et al*. (1978) observed cell cycles under the light microscope without embryo exposure to laser beams. Laser exposure to embryos caused developmental delays of ∼six hours by E3.5 as discussed in (Yang, 2025b), which may explain the differences in observed cell cycle durations. The differences between my findings and those reported by Bischoff *et al*. (2008) could stem from differences in experimental conditions, such as video conditions and the transgenic strains used. Additionally, cell cycle length variations increased as embryos grew in this study, especially after the 32-cell stage (Gn5). While the detailed causes of these variations are beyond the scope of the present work, inherent or acquired differences in cell differentiated cell lineages and cell spatial properties may underscore the variations, as discussed below.

### 3.5 Cell Cycle Length and Cell Fate Decision

Bischoff *et al*. (2008) reported longer cell cycles in inside cells than outside cells once their positions were determined. My study drew on that, showing cells located relatively inside/closer to the embryo centre consistently outlasted those of outside cells through Gn1 to Gn6, until the early-mid stage of Gn7. This indicates that early “differentiated” TE cells proliferate faster than pluripotent-like ICM cells, highlighting the dynamic spatial and temporal trends in cell cycles. However, in Gn7-Gn8, P-TE cells began to show shorter cycles while Mu-TE cells had longer cycles (18 or over 22.5 hours), with pairwise sisters loosely surrounded by neighbours having shorter cell cycles. Supports can be seen from previous studies that showed new cell cycles of TE cells distinct from the ICM, with mitoses restricted to P-TE cells around the implantation stages (Gardner, 2000; Gardner and Davies, 2002). My results also showed that Epi and PrE precursors had similar cell cycle lengths in Gn6, but in Gn7, PrE-like cells exhibited varied cell cycles, with cells in the ICM margin zone appearing with shorter cell cycles than the surface and deep ICM. My analyses between pairwise daughters at 4-, 8-and 16-cell stages showed that the daughters of short-cycled blastomeres internalised more in the 4^th^ division round, forming more Mu-TE (near P-TE) cells whose sisters usually were early-dividing marginal ICM cells. These findings extended our understanding of the relationships between cell cycles and cell fate decisions, which was traditionally thought that differentiated cells had longer cell cycles (Copp, 1978; Pauklin and Vallier, 2013). My study suggests that during early embryo development, TE, ICM, PrE, Epi and their sublineages experience dynamic transitional changes in their cell cycles and cell fates, driven by cell origins, positions and the inheritance of their historical spatiotemporal features.

### 3.6 Interplay between Cell Division Order, Cycle, Position, and Lineage Specification

Cell cycle history analyses in this study showed a novel non-linear alternative yet specific shifts in cell cycle lengths across generations, based on odd-or even-numbered cycle categories, embryo stages and pairwise sister positions and cell types, including mother, daughter, sibling, and cousin cells. This indicates that cell cycle durations are stage-dependent, cycle history-associated and position-driven. Furthermore, some 4-cell descendants from faster-dividing 2-cell blastomeres exhibited extensively long cell cycles in Gn7, particularly in the Mu-TE cells. These findings further suggest that potential cell property rearrangements in each division round and their subsequent inheritance may affect cell cycle regulation and cell allocation to ICM/TE across cell generations. This was evidenced by my analyses where short-cycle 2-, 4-and 8-cell embryonic cells internalised descendants earlier and produced more marginal ICM (PrE) and more Mu-TE (near P-TE) by E4.5. This study supports the work by Krawczyk *et al*. (2021) showing individual 2-cell blastomeres or quart droplets from 4-cell embryos exhibit distinct capacities of forming subsets of TE, PrE and Epi.

This present study revealed that cell cycle and division orders interacted dynamically and non-linearly over time, challenging the idea of no decisive link between these two factors in early mouse embryos concluded by Kelly *et al*. (1978). My study showed that cell histories of division order, origins and cycles generally modulated division order and cell cycle lengths of descendants, with cell positions also playing key roles within each generation. Cells with fewer surrounding neighbours (outside cells at the morula stage, Mu-TE and marginal ICM cells) consistently divided earlier and showed shorter cell cycles than cells with more neighbours (those near the embryo centre at the morula stage, deep ICM cells, and P-TE cells), especially evident by the late 32-and 64-cell stages. At the late 32-and 64-cell stages, cells with few neighbours, such as Mu-TE (near P-TE) and marginal ICM (potential PrE) tended to have longer cell cycles, while cells with more surrounded cells (P-TE and deep ICM) started to show relatively shorter cell cycles, potentially due to functionally matured P-TE near Mu-TE and PrE cells. My findings extended previous work by Forsyth *et al*. (2021) showing that cell neighbours of ICM, Mu-and P-TE cells changed over time by illustrating how these changes in neighbouring packing influence cell cycles.

### 3.7 Cell Cycle Phases and Developmental Events

This study showed that cells undergoing initiation of compaction, cavitation and hatching events were in the early phases of cell cycles. Although the detailed cell cycle phases, such as G1, S, M and G2, are beyond the scope of technical approaches used here, my findings provide a general understanding of the coordination between specific phases of cell cycles and the distinct phases of embryo morphological events, implying the potential regulatory roles of cell cycle and their phases in embryo development. Traditionally, the developmental timing of morphological events is usually defined and assessed by developmental days in both mouse and human embryos (Xue *et al*., 2013; Wong *et al*., 2015). However, there has been limited analysis of the relationships between cell cycle phases and morphological events. My study extends the temporal framework for assessing embryo development and also underscores the need for further investigation into the detailed mechanisms of cell cycle phases coupling with embryo morphogenesis.

## 4 Conclusions

The study reveals intrinsic spatial and temporal relationships between cell positions, division orders, cell cycle dynamics, cell lineages and various morphological events during pre-and early peri-implantation stages. It highlights the impact of historical spatiotemporal cues (such as cell positions) on driving cell division orders and cell cycles, consequently directing asymmetric cell lineage specification of individual ancestor blastomeres. Pairwise sisters leverage these spatiotemporal cues to guide cell fate decisions, and balance cell fate asymmetry and the integrated developmental symmetry. These findings enhance our understanding of historical developmental trajectories, and shed light on how embryonic cells balance heterogeneous properties across pairwise blastomeres while maintaining homogenous features across generations via inheritance and how different cell populations interdependently coordinate their different differentiation statuses as embryos develop. These insights have potential applications in stem cell study, regenerative medicine and ART fields. Future work is recommended to validate these findings by exploring underscoring molecular mechanisms of inheritance and differentiation of spatiotemporal cues directing embryo morphogenesis and cell lineage specification.

## 5 Study Limitations and Prospects for Advancing Research and Future Directions

In this work, embryonic cell tracking of cell position, division, cycle, origin and differentiation relied on H2B-GFP marker, which offers novel insights into spatiotemporal dynamics of cell lineage origin and differentiation. To refine these insights and explore their underlying mechanisms, future work should focus on developing and integrating advanced molecular methods for spatiotemporal live cell tracking and single-cell multi-omics analysis. Additionally, combining time-lapse imaging with embryo culture and molecular techniques to capture complete peri-and post-implantation development would further enhance the depth and accuracy of these analyses.

## 6 Materials and Methods

### 6.1 Ethical Approval and Regulations for Animal Care

For this work, no new animals were used. Embryos and embryonic videos from mice reported in (Yang, 2025b) were reused for different data analyses to address new research questions and hypothesis in the present study, complying with the 3Rs principles (Replacement, Reduction and Refinement). Ethical approval and animal care regulations for the original animals used are detailed in (Yang, 2025b).

### 6.2 Experimental Design

#### 6.2.1 Workflow and Parameters for Investigation

The study designs for this work, which align with the hypothesis and objectives, are presented in (Figure 15). Independent variables (IVs) primarily related to distinct developmental stages, cell positions, cell division orders, and cell durations, while the dependent variables (DVs) mainly included cell lineages or cell populations. Cell positions, cell division orders, and cell durations were also assigned as IVs or DVs for specific analysis aims, depending on the research questions raised. Notably, each group was designated as an internal control for the others within its respective set, addressing confounding variables potentially influencing embryo development and research outcomes, excluding IVs and DVs. Furthermore, short videos served as a control group for long videos in the context of temporal factor analysis. The detailed parameters to be measured and analysed are outlined in (Figure 15).

**Figure 15.**
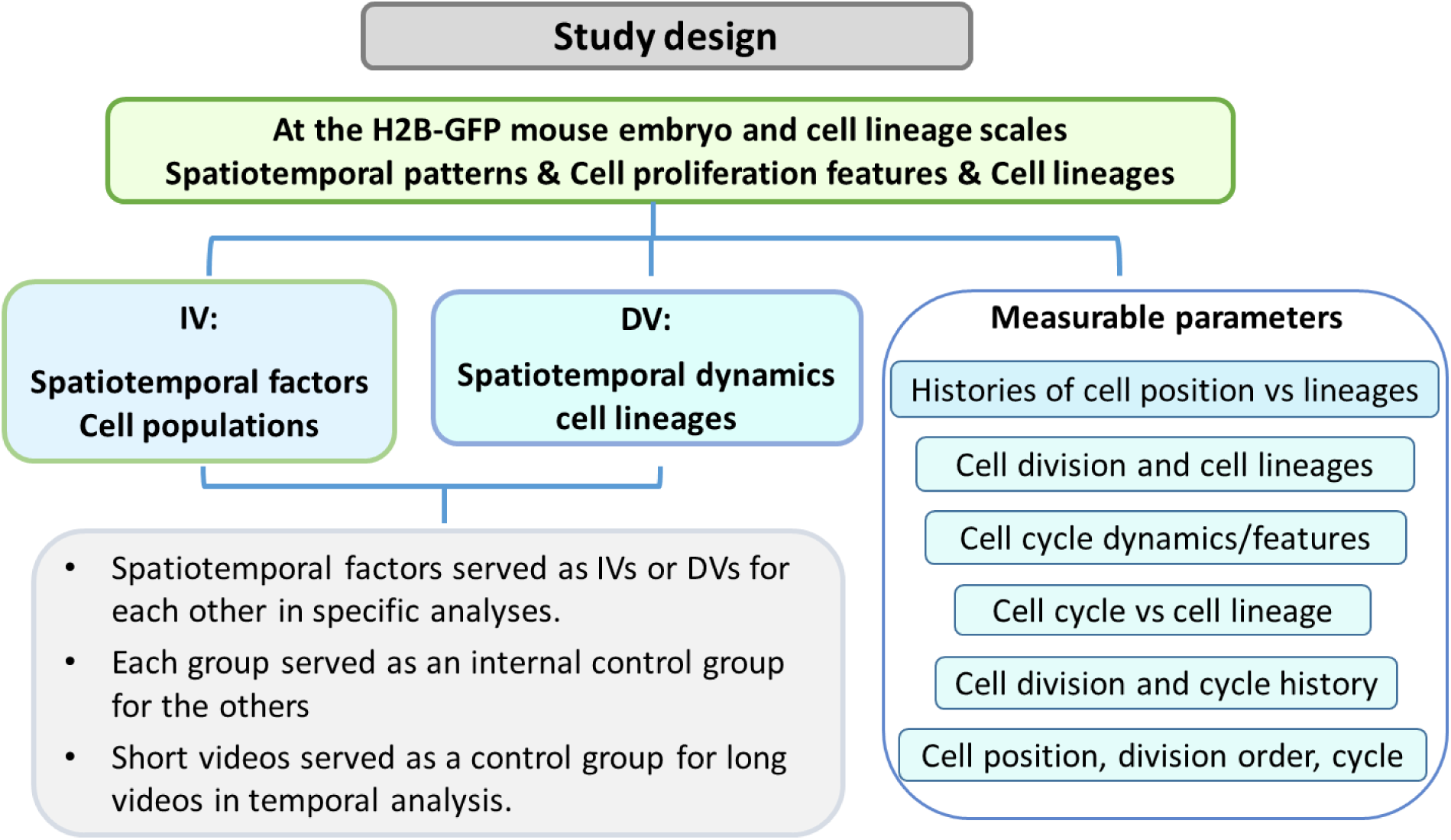
Experimental designs of mouse embryo study. The top box in light green/yellow displays the major scales and the major factors investigated in this study. The left blue boxes in the middle depict IVs (independent variables) and DVs (dependent variables), followed by further explanation for IVs, DVs and control group settings in the grey box. The right box shows the main parameters to be measured and analysed in this work.

#### 6.2.2 Inclusive and Exclusive Criteria

Based on the typical development of mouse embryos as reviewed in (Yang, 2025a) and in line with my study aims, the four inclusive criteria of embryos were defiend as previously described in (Yang, 2025b), unless stated otherwise. These inclusive criteria were repeated here for easy reference: “(1) embryos survived and developed to the pre-or peri-implantation stages; (2) embryos displayed the stage-characterised morphology landmarks such as compaction and cavitation in the time course of interest as described in (Yang, 2025a); (3) late blastocysts exhibited proper proportions of lineages to be examined, and that was assessed through both live-cell and fixed-cell images; and (4) transgenic embryos showed the visible and trackable intensity of targeted vector.” To thoroughly investigate historical spatiotemporal patterns of early embryo development and control any confounding factor that might influence the parameters designed to be measured, the exclusive criteria included: (1) embryos with dying cells before the 32-cell stage were excluded (dying cells were identified by failure to divide and having fragmented nuclei under a fluorescence microscope); (2) embryos arrested at the 2-, or 4-cell stages, and failing to progress into subsequent developmental events within the expected time frame were exempted from long time-frame analyses, refer to (Yang, 2025a) for a reviewed embryo developmental timeline; (3) embryos that drifted dramatically under the microscope were not included for long-time cell track; and (4) embryos that had collapsed to a significant extent, marked by loss of cavity and having irregular embryo shape under a microscope, were excluded from the match between live cell images and their corresponding fixed-cell images. These criteria were consistently applied to all procedures in the embryo study, including embryo collection and image analyses.

#### 6.2.3 Sample Size

The sample size was calculated using a large estimated effect size (Cohen’s d ≈ 1.78) for cell cycle lengths between outside and inside cells. Accordingly, an initial sample size of six embryos per group was deemed to be sufficient to detect meaningful trends in the aimed DVs for long-term live imaging of embryos. The final sample size was increased to 10 embryos (unless otherwise noted) to enhance robustness and ensure more reliable detection of meaningful differences between the analysed groups, facilitated by additional access to the Bioimaging Facility (UoM) for cell tracking. For short movies, a minimum of three to four embryos was considered sufficient due to the reduced variability of each embryo development over time and the less complex nature of measurements.

### 6.3 Laboratory Work Procedures

In this study, embryos and embryonic videos from mice reported in (Yang, 2025b) were reused for different data analyses to align the study design. The procedural details of the laboratory work, key resources (such as chemicals and apparatus) and summarised tables are as previously described (Yang, 2025b).

### 6.4 Imaging analysis and Data collection

#### 6.4.1 General Image Analysis

Acquired ND2 files obtained from the imaging system were converted to IMS files via IMARIS software for cell tracking as described in (Yang, 2025b). Inkscape was used to assist with matching embryonic cell positions by adjusting transparency levels and juxtaposing live embryo images with their corresponding fixed embryo images. The resulting matches were then compared with my manual-identified matches to enhance accuracy. The software/tools used for this are summarised in (Table 2). Following this, manual corrections were made for missed or overlapped objectives using IMARIS. Embryonic cells were tracked at each time point in each movie (refer to Section 6.4.4 for details of cell tracking). Cells that were dying or had too weak signals hindering accurate tracking were marked and excluded. All other embryonic cells were successively tracked, except for the excluded embryos based on the criteria described in Section 6.2.2. To maintain the consistency of the tracking process, I double-checked the cell tracks three times every six months. Afterwards, parameters aimed for analyses, such as cell cycle, and cell positions, were either manually collected from cell tracking trees and movies per se or automatically extracted from IMARIS.

**Table 2.**
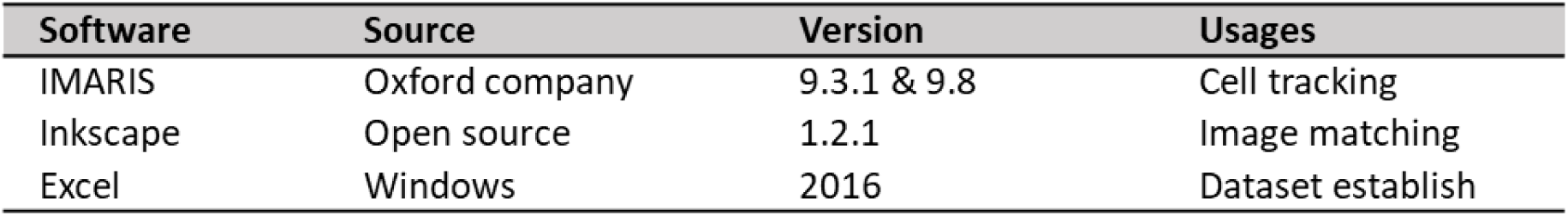
Software for image analyses. This table contains the main software used for imaging, processing and data collection in mouse embryo work.

#### 6.4.2 Counting embryonic cells

Embryonic cell numbers count are as previously described (Yang, 2025b).

#### 6.4.3 Monitoring embryo morphological changes

Embryo morphological events were measured as described in (Yang, 2025b) for *in-vitro* cultured embryos.

#### 6.4.4 Cell tracking and matching in 4D time-lapse videos of H2B-GFP mouse embryos

During tracking, time points corresponding to cell divisions were identified based on the morphology, numbers, and intensity of H2B-GFP-visualised nuclei (Figure 16A,B). Specifically, the time point when clear separation of two sets of H2B-GFP-labeled chromatin was detected indicated the start of the following cell generation. Integrating different strategies of image processing and both retrospective and prospective cell tracking approaches (Figure 16B), I tracked each single embryonic cell to the fullest extent when feasible. To reconstruct various cell lineage histories within the long 4D films, I cross-referenced and matched two distinct sets of images: live embryo images from the 2-cell stage to the late blastocyst stage, and fixed embryo images consisting of the same late blastocysts where cells were immunostained by Sox2 and Gata4 indicative of Epi and PrE, respectively, along with Hoechst-stained nuclei (Figure 16C). Embryo matching was based on comparing embryo morphology, total cell number, and any specific features which could serve as a landmark within embryos. Following this, embryonic cells were recognised and paired based on their positions, shapes, adjacency, and any identifiable recognition landmarks. These visual and manual matching procedures were cross-validated using Inkscape software via registering two sets of images, as described in Section 6.4.1.

**Figure 16.**
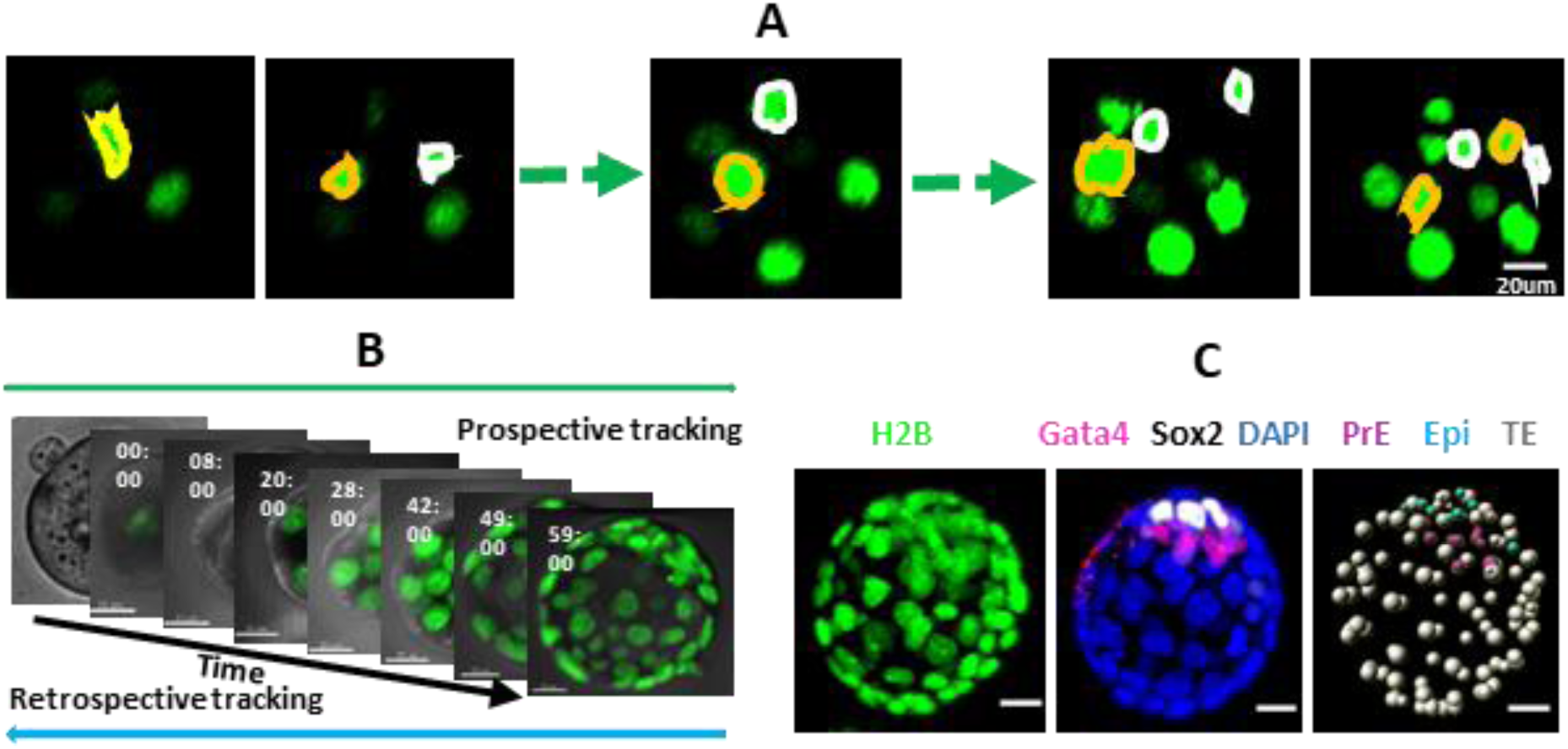
Embryonic cell tracking, matching and lineage identification in long videos of H2B-GFP mouse embryos. **(A)** Cell recognition over time with highlighting cell nuclei features during cell divisions; the yellow and white circles represent two distinct mother cells and their corresponding daughters. **(B)** Retrospective (blue arrow) and prospective (green arrow) refer to cell tracking directions from two cells to over 100 cells. The black arrow indicates consecutive timing. **(C)** Embryonic cell match among H2B live cells, immunostained cells and tracking spots in IMARIS between live and immunostained images, with green, white, red and dark blue representing H2B, Sox2, Gata4 and DAPI (Hoechst) florescence-positive cells, respectively. Scale bar in (A-C): 20 µm.

#### 6.4.5 Identifying and categorising cell positions within H2B-GFP mouse embryos

Unless otherwise specified, embryonic cell positions were determined based on fluorescence-labelled nuclei, identified using the IMARIS spot algorithm. The 3D coordinates of cell positions in long-term embryo videos or fixed snapshot images were extracted from IMARIS for further analysis. To classify cell positions within embryos, distances from each cell and the geometric centre of the embryo were measured. Cell positions were then ranked based on absolute distances or grouped into equal ranks for specific analysis. For the ranking, TE cell positions were categorised along a distance gradient from P-TE apex to Mu-TE apex. ICM cell positions were ranked from deep to surface layers in equal distance increments, with a negative notation distinguishing ICM ranks from TE cells.

#### 6.4.6 Cell cycle duration measurement

Cell cycle lengths were investigated for cells that underwent at least two consecutive and discernible cell divisions, with the duration measured from the start of one cell division to the start of the next division. Cells that experienced fewer than two successive rounds of divisions or displayed abnormal morphological changes were excluded from the cell cycle length analysis.

### 6.5 Statistical Analysis and Data Visualisation

This section was conducted as previously described (Yang, 2025b).

## 7 Acknowledgements

This work was from my self-funded research at the University of Manchester. I thank the colleagues at the University of Manchester for their comments on this work. My thanks also go to the core facilities at the university, especially the Bioimaging Facility and Animal Facility for their invaluable support and expertise throughout the research process.

## Appendix A

### Specific 24-hour time patterns of embryonic cell divisions during pre-and peri-implantation stages

While collecting freshly flushed embryos in (Yang, 2025b), I observed specific timings of cell divisions across various developmental stages; the observed divisions in the 1-cell, 2-cell, 4-cell, 8-cell, 16-cell, 32-cell, and 64-cell embryos were termed here the first through seventh embryonic division rounds (denoted as the 1^st^, 2^nd^, 3^rd^, 4^th^, 5^th^, 6^th^, and 7^th^, respectively). As such, I investigated whether early embryonic cell divisions possessed consistent 24-hour timing patterns. Critical evolution of animal events used in (Yang, 2025b) provided a uniform baseline across all embryos for determining 24-hour time of cell divisions (assessed in Appendix B). The results revealed that *in-vivo* embryonic cells showed odd division rounds (3^rd^, 4^th^, and 5^th^) mostly during the day, and even division rounds (2^nd^ and 4^th^) with the 1^st^ round clustering on night hours (Figure A1). Direct observation of cell division timings at the 64-cell stage was challenging due to difficulties in detecting cell division, but extrapolation from observed earlier-stage divisions suggested that 64-128 cell divisions could occur after 18:00 (Figure A1Ac, dashed lines). To confirm this prediction, I examined short H2B-GFP videos of six 32-cell embryos for division timings in the 6^th^ and 7^th^ rounds. The 6^th^ division round spanned from 18:00 to 22:00, and the 7^th^ round was from 07:00 to 12:00, with both rounds presenting a wider range than previous division rounds (Figure A1B). However, not all of these cells could be definitively attributed to the 7^th^ generation due to overlapping division timings between the 6^th^ and 7^th^ rounds, and between the 7^th^ and 8^th^ generations.

Longitudinal data of H2B-GFP embryos from E1.5 to E4.5/E4.75 (Figure A1C), with sample sizes summarised in (Figure A1Da), confirmed the consistent patterns of 24-hour times of cell divisions. The median timings for each division round, with the 25^th^ and 75^th^ percentiles in parentheses, were as follows: the 1^st^ round at 23:50 (22:30-01:00), the 2^nd^ round at 20:10 (18:00-22:00), the 3^rd^ round at 07:20 (06:20-10:39), the 4^th^ round at 18:00 (17:30-19:14), 5^th^ round at 06:00 (04:40-09:30), the 6^th^ round at 17:00 (15:00-20:40), and the 7^th^ round at 10:00 (07:00-12:00). The 8^th^ round, measured in only 11 cells within three embryos, ranged from 09:00 to 12:30. Interestingly, odd division rounds (3^rd^, 5^th^, 7^th^) primarily occurred in the morning (06:00-10:00), while even rounds (2^nd^, 4^th^, 6^th^) occurred at night (18:00-20:00) (Figure A1Db).

As the latest three rounds exhibited extended durations with skewed distributions, I examined the relationships between the earliest and latest two to three hours of division times and cell positions. In six embryos, cells dividing earliest (12:00-14:00) were 78.57% TE cells located at the boundary between P-TE and Mu-TE (termed Ma-TE), 7% Mu-TE, 7.43% P-TE and 7% ICM, while 48 cells dividing latest (0:00-3:00 on the following day) were 62.5% Mu-TE and 37.5% ICM (n=42 cells) (Figure A1Ea). In the 7^th^ round, the earliest-dividing cells (2:00-4:00) were 65% Ma-TE, 18.3% P-TE and 16.7% ICM (n=60 cells), while the cells dividing latest (11:00-13:00) were 72% Mu-TE, 27% ICM and 1% P-TE cells (Figure A1Eb). In the 8^th^ round, although not all generation cells were examined due to the limited video durations, the earliest dividing cells (8:00-11:00) were 50% P-TE, 33.3% ICM and 16.7% Ma-TE (n=12 cells) (Figure A1Ec).

**Figure A1.**
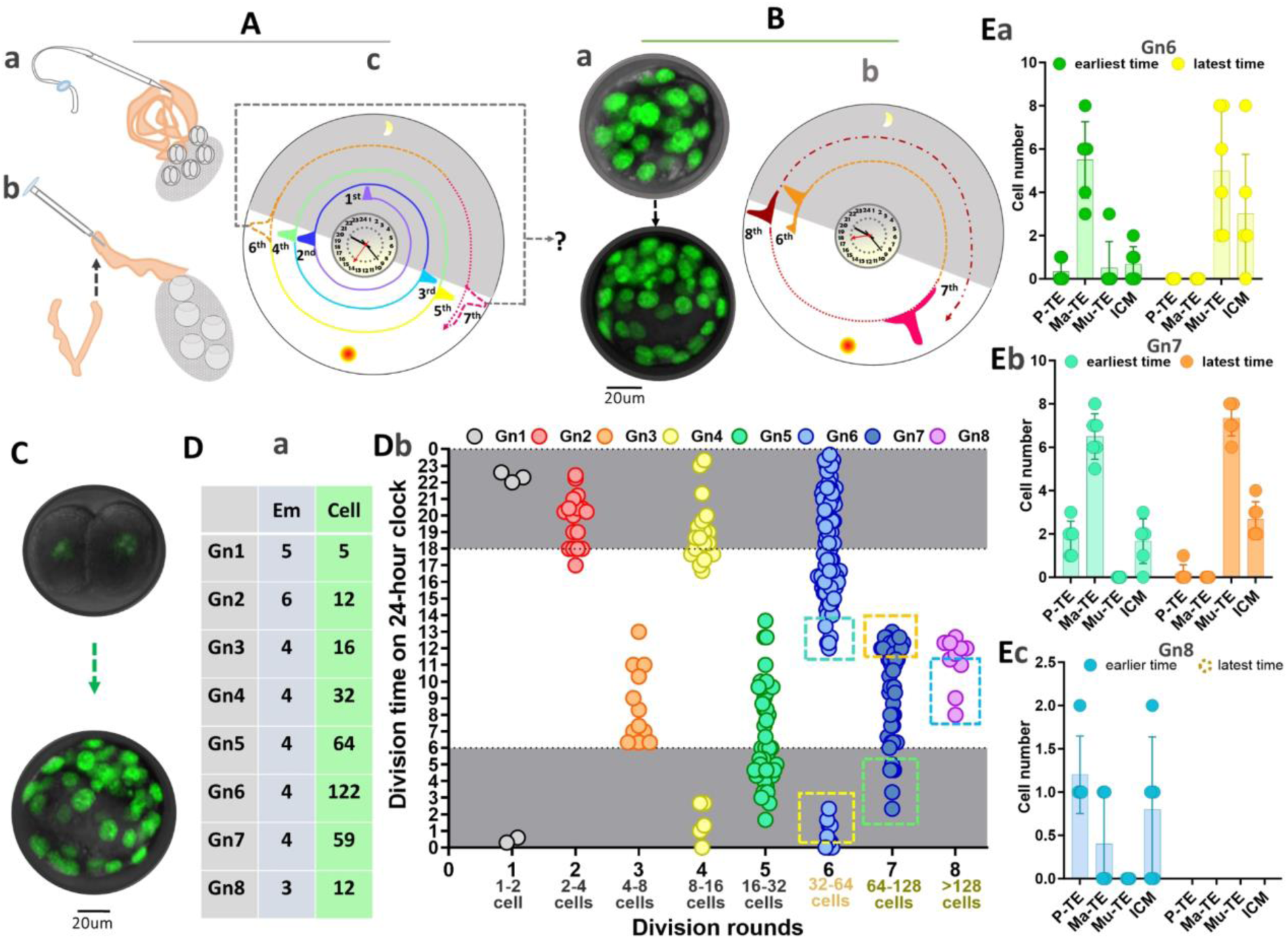
The 24-hour time patterns of embryonic cell divisions. **(Aa,b)** Observations of cell division time during embryo collection; concluded from the work in (Yang, 2025b). **(Ac)** General 24-hour timings of *in-vivo* cell divisions, with the 1^st^ to 5^th^ rounds colour-coded in purple, dark blue, blue, green, yellow, 6^th^ and 7^th^ in orange and red dashed curves, respectively. **(Ba,b)** General 24-hour timings of cell divisions in the 6^th^ (orange), 7^th^ (red) and 8^th^ (dark red) rounds within short videos of H2B-GFP embryos. **(C,D)** Cell division timings in 4D long live embryo videos, with sample size summarised in (Da) and detailed cyclic patterns of specific 24-hour division timing illustrated in (Db) (white refers to daytime and grey indicates nighttime). Cell generations/division rounds were colour-coded as shown in (Db), with pairs of green/blue and yellow dashed boxes in Gn6 and Gn7/Gn8 indicating cells dividing at the earliest and latest 24-hour clock in each round, respectively. **(Ea-c)** Cell numbers with SD error bars of various cell types dividing in the earliest (green/blue) and latest (yellow/orange) 24-hour clock in Gn6, Gn7 and Gn8, respectively, corresponding to (Db). The scale bar in Panels B and C: 20 µm.

## Appendix B

### Conceptual framework for embryonic division timing and progression

To maintain consistency and clarity in understanding the observed temporal events in early embryos, I developed a conceptual framework. Specifically, I introduced the term ‘embryonic progression period’ to describe observed embryonic division rounds and intervals following each round. Within each period, I defined ‘embryonic division (round) durations’ as the time length characterised by continuous cell divisions in each round, and ‘embryonic division intervals’ as the time window where no cells underwent mitosis. These concepts were built to distinguish embryonic progression, featuring collective cell division events at the population level, from individual embryonic cell life cycles.

## Appendix C

### Conceptual frameworks for cell cycle progression

To systematically describe the features of cell cycle lengths in each single embryonic cell, I first formulated a conceptual framework outlining relations between cell generations and cell cycle rounds. Cell generations were referred to cohorts of cells stemming from the same cell origin/ancestor. Within a representative of cell cycle trees displayed in (Main thesis text, Figure 4.9, Panel A), each level represented a generation (Gn) of the cell cycle, with the top level being the first generation, and subsequent levels representing the second (Gn2) through the 8^th^ generations (Gn8). Corresponding to these generations. Embryonic cell cycle periods, *i.e.*, embryonic progression periods, referred to as a time window during which the first cell in their generation starts to divide until the first cell in the next generation starts to divide, termed 1^st^, 2^nd^, 3^rd^, 4^th^, 5^th^, 6^th^, 7^th^, 8^th^. Cell generations and cell cycle periods signified the corresponding respective 1-cell, 2-cell, 4-cell, 8-cell, 16-cell, 32-cell, 64-cell and 128-cell stages. At the 1-cell stage, cell cycle lengths were computed based on visual observation of freshly flushed embryos which were undertaking division.

